# A negatively-charged region within carboxy-terminal domain maintains proper CTCF DNA binding

**DOI:** 10.1101/2024.04.14.587166

**Authors:** Lian Liu, Yuanxiao Tang, Yan Zhang, Qiang Wu

## Abstract

As an essential regulator of higher-order chromatin structures, CTCF is a highly-conserved DNA binding protein with a central DNA-binding domain of 11 tandem zinc fingers, which are flanked by amino (N-) and carboxy (C-) terminal domains of intrinsically disordered regions. Here we report that CRISPR deletion of the entire C-terminal domain of alternating charge blocks decreases CTCF DNA binding but deletion of the C-terminal fragment of 116 amino acids results in increased CTCF DNA binding and aberrant gene regulation. Through a series of genetic targeting experiments, in conjunction with EMSA, 4C, qPCR, ChIP-seq, and ATAC-seq, we uncovered a negatively-charged region (NCR) responsible for weakening CTCF DNA binding and chromatin accessibility. AlphaFold prediction suggests an autoinhibitory mechanism of CTCF via NCR as a flexible DNA mimic domain, possibly competing with DNA binding for the positively-charged ZF surface area. Thus, the unstructured C-terminal domain plays an intricate role in maintaining proper CTCF-DNA interactions and 3D genome organization.

## INTRODUCTION

CCCTC-binding factor (CTCF) is a principal architectural protein for the construction of 3D genomes and is highly conserved in bilateria.^1–5^ Together with the cohesin complex, CTCF mediates the formation of long-distance chromatin loops between distant sites, known as CBS (CTCF binding site) elements, through an ATP-dependent active process known as “loop extrusion”, leading to higher-order chromatin structures such as TADs.^6–13^ Interestingly, CTCF/cohesin-mediated chromatin loops are preferentially formed between pairs of CBS elements in a forward-reverse convergent orientation.^14–17^ In particular, topological chromatin loops are formed between tandem-arrayed CBS elements via cohesin-mediated dynamic loop extrusion, leading to balanced promoter choice.^18–20^ The dynamic cohesin loop extrusion and its asymmetric blocking by oriented CTCF binding on numerous CBS elements distributed throughout mammalian genomes constitute a general principle in 3D genome organization and play an important role in gene regulation. Finally, other DNA-binding zinc-finger proteins such as YY1, MAZ, PATZ1, and ZNF263 may collaborate with CTCF to form long-distance chromatin contacts.^21,22^

The clustered protocadherin (*cPcdh*) genes are an excellent model to investigate the relationships between CTCF/cohesin-mediated chromatin looping and gene expression programs. The 53 highly-similar human *cPCDH* genes are organized into three tandemly-linked clusters of *PCDHα*, *PCDHβ*, and *PCDHγ*, spanning a large region of ∼1M bps genomic DNA.^23^ The *PCDHα* gene cluster comprises an upstream region of 15 variable exons and a downstream region with 3 constant exons. Similarly, *PCDH*γ comprises an upstream region of 22 variable exons and a downstream region of 3 constant exons. Each variable exon is separately spliced to the respective set of 3 constant exons within the *PCDH α* or *γ* gene cluster. By contrast, the *PCDHβ* gene cluster comprises only 16 variable exons with no constant exons.

Similar to the intriguing *Dscam* gene for generating enormous diversity of cell-recognition codes in fly, the *cPCDH* genes generate an exquisite diversity for neuronal self-avoidance and nonself discrimination in vertebrates.^24,25^ Different from competitive RNA pairing-mediated mutually exclusive splicing mechanism for *Dscam*, the *cPCDH* diversity is generated by a combination of balanced promoter choice and *cis*-alternative splicing determined by CTCF-directed DNA looping.^14,26^ In this complicated 3D genome configuration, CTCF directionally binds to tandem arrays of oriented CBS elements associated with *Pcdh* variable promoters and super-enhancers. CTCF-mediated chromatin loops are then formed between pairs of convergent forward-reverse CBS elements. For example, in the *PCDHα* gene cluster, there are two forward CBS elements flanking each of the 13 alternate variable exons and two reverse CBS elements flanking the *HS5-1* enhancer.^27^ A “double-clamping” chromatin interaction between these convergent pairs of CBS elements determines the *cPCDH* promoter choice.^14^ In summary, CTCF/cohesin-mediated loop extrusion bridges remote super-enhancers in close contact with target variable promoters to form long-distance chromatin loops and this looping process is essential for establishing proper expression patterns of the c*PCDH* genes in the brain.^14,17–19,28–30^

CTCF contains a central domain of 11 zinc fingers (ZFs) organized in a tandem array flanked by intrinsically disordered regions of the N-terminal domain (NTD) and C-terminal domain (CTD) (Figure 1A).^1,31,32^ The central domain of CTCF binds to DNA directly through ZFs 3-7 and ZFs 9-11.^17,33–35^ Recently, several lines of evidence suggest that ZF1 and ZF2 recognize base pairs downstream of the CTCF core motif.^36–40^ Remarkably, CTCF also interacts with RNA to mediate chromatin loop formation and to regulate gene expression pattern.^41,42^ Finally, the intrinsically disordered NTD, but not CTD, of CTCF interacts with cohesin complex to anchor chromatin loops between distant DNA elements.^4,12,43–46^ Here by a combination of a series of genetic deletions, in conjunction with chromosome conformation capture and gene expression analyses, we found that a negatively-charged region (NCR) within the disordered CTD is important for proper CTCF DNA binding, higher-order chromatin organization, and gene regulation.

**Figure 1.**
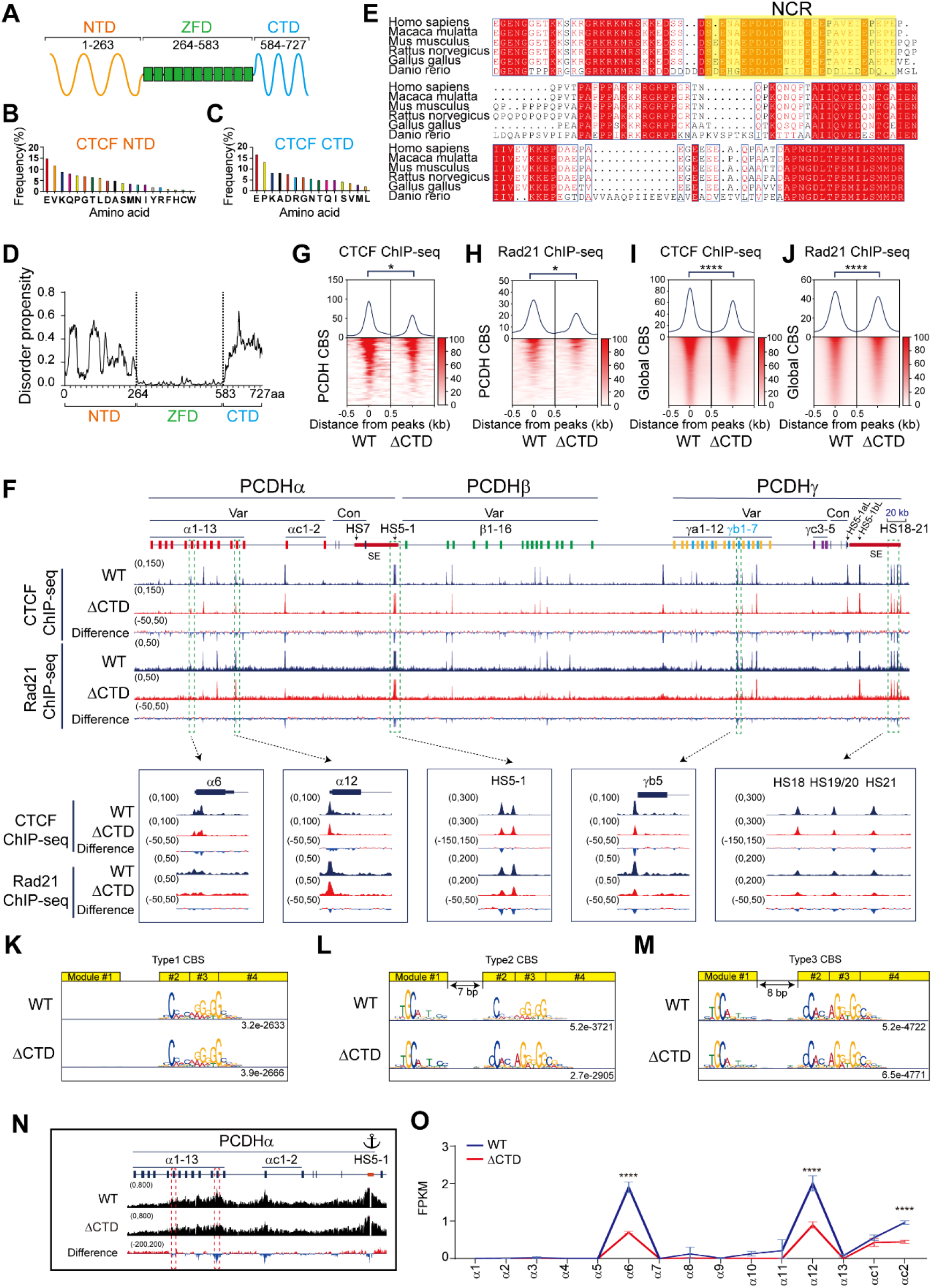
Deletion of CTCF C-terminal domain results in decreased CTCF binding and gene dysregulation. (A) Schematic of the CTCF zinc finger domain (ZFD) and the flanking N-terminal domain (NTD) and C-terminal domain (CTD). (B and C) Amino acid composition of CTCF NTD (B) and CTD (C). (D) Disordered propensity by a computer program indicates that NTD and CTD are intrinsically disorder region (IDR). (E) Multiple sequence alignment of CTD of the vertebrate CTCF proteins. The negatively-charged region (NCR) is highlighted in a yellow background. (F) CTCF and Rad21 ChIP-seq peaks at the human *cPCDH* gene complex. The human *cPCDH* locus comprises three tandemly-linked gene clusters: *PCDHα*, *PCDHβ*, and *PCDHγ.* The *PCDHα* and *PCDHγ* clusters each consist of a variable region with multiple highly-similar and unusually-large exons and a constant region with three small exons. Each variable exon is separately *cis*-spliced to a single set of cluster-specific constant exons. The *PCDHβ* cluster contains 16 variable exons but with no constant exon. These three clusters form a superTAD with two subTADs: *PCDHα* and *PCDHβγ*. Each subTAD has its own downstream super-enhancer. Var, variable; Con, constant; HS, hyper-sensitive site; SE, super-enhancer. (G and H) Heatmaps of CTCF (G) and Rad21 (H) normalized signals at CBS elements in the *cPCDH* locus. Student’s *t* test, \**p* < 0.05. (I and J) Heatmaps of CTCF (I) and Rad21 (J) normalized signals at genome-wide CBS elements. Student’s *t* test, \*\*\*\**p* < 0.0001. (K-M) Three types of CTCF motifs in WT and ΔCTD cells. (N) QHR-4C interaction profiles of the *PCDHα* gene cluster using *HS5-1* as an anchor. (O) RNA-seq shows decreased expression levels of *PCDHα6, PCDHα12,* and *PCDHαc2* upon deletion of CTD. FPKM, fragments per kilobase of exon per million fragments mapped. Data are presented as mean ± SD; Student’s *t* test, \*\*\*\**p* < 0.0001. See also Figures S1, S2, and S3.

## RESULTS

### Deletion of CTCF CTD results in decreased DNA binding

We analyzed the amino acid (AA) composition of CTCF NTD and CTD and found that the NTD of CTCF contains all of the 20 types of amino acids while the CTD of CTCF contains only 15 AA types, suggesting that CTCF CTD has an amino acid compositional bias and is a low complexity region (Figure 1B and 1C). Computational analyses suggest that both NTD and CTD regions have a high intrinsically disordered score, especially the CTD region (Figure 1D).^47^ In addition, the CTD region is highly conserved in vertebrates (Figure 1E). Owning to the lethality of CTCF deletion in mice or in cultured cells,^48,49^ we tried to delete the CTCF NTD or CTD in HEC-1-B^17^ and found that deletion of NTD, but not CTD, is lethal in cultured cells. Specifically, we screened 254 single-cell clones for deletion of NTD, and could not find a single homozygous cell clone. Therefore, we focused our genetic dissection on CTCF CTD.

We screened for CTD-deletion clones by CRISPR DNA-fragment editing programed with dual sgRNAs and a donor construct containing FLAG sequences for tagging^50,51^ and obtained two single-cell clones with precise deletion of the CTCF CTD (ΔCTD) (Figure S1A-D). We performed CTCF ChIP-seq with these two ΔCTD clones as well as with WT clones as a control (Figure S1B) and found that almost every CTCF peak within the three *PCDH* gene clusters is decreased upon CTD deletion, suggesting that CTD has an important role in CTCF binding to DNA (Figures 1F and S2A, S2B). Aggregated peak analysis showed that there is a significant decrease of CTCF binding at the *PCDH* CBS elements (Figure 1G). DNA-bound CTCF anchors cohesin complex via its NTD but not CTD.^4,12,43–46^ To this end, we performed ChIP-seq experiments with a specific antibody against Rad21, a cohesin subunit, and found that cohesin is colocalized with CTD-deleted CTCF, suggesting that CTD-deleted CTCF is still able to anchor cohesin at the *cPCDH* locus (Figures 1F and S2A). However, there is a significant decrease of cohesin enrichments upon deletion of CTCF CTD (Figures 1F, 1H and S2A, S2C). We then analyzed genome-wide CTCF and cohesin enrichments and found that both CTCF and cohesin enrichments are significantly decreased upon CTD deletion (Figure 1I and 1J). However, computational analyses revealed no alternation of CTCF motifs of all three types of CBS elements upon CTD deletion, suggesting that CTD does not alter the DNA binding specificity of the central ZF domain (Figure 1K-1M), despite the fact that CTCF enrichments are decreased for all three types of CBS elements (Figure S2D-S2F).

We next performed quantitative high-resolution circularized chromosome conformation capture experiments (QHR-4C, see methods)^19^ with the *HS5-1* enhancer as an anchor and found that there is a significant decrease of long-distance chromatin interactions between the *HS5-1* enhancer and its target promoters (Figure 1N). Finally, we performed RNA-seq experiments and found that, consistent with decreased chromatin interactions between enhancers and promoters, there is a significant decrease of expression levels of members of the *PCDHα* gene cluster upon CTCF CTD deletion (Figure 1O).

### Deletion of CTCF CTD affects gene regulation

We next analyzed the RNA-seq data using DESeq2 with adjusted *P* value < 0.05 and log2FC (fold change) >1 as cutoffs. We found 150 up-regulated genes (Table S1, ingenuity pathway analysis IPA, Figure S3A) with mean log2FC of 1.70 and 207 down-regulated genes (Table S2, Figure S3B) with mean log2FC of −1.84 (Figure S3C). We also found that the down-regulated genes are closer to CBS elements than the up-regulated genes (Figure S3D).

### Deletion of C-terminal 116 AAs leads to increased CTCF binding

The CTD of CTCF contains an internal RNA-binding region (RBR) and a downstream region of 116 amino acids.^42,52^ To investigate its function, we generated targeted deletion by screening single-cell CRISPR clones using CRISPR DNA-fragment editing with Cas9 programmed by dual sgRNAs.^17,50^ We obtained two clones with deletion of the C-terminal 116 AAs (ΔCT116) (Figure S1E-G). We performed CTCF ChIP-seq experiments and found, remarkably, that there is a significant increase of CTCF enrichments in the *cPCDH* gene complex upon deletion of the C-terminal 116 AAs (Figures 2A, 2B and S4A, S4B). In addition, we performed Rad21 ChIP-seq experiments and found that there is a significant increase of cohesin enrichments at the *cPCDH* locus (Figures 2A, 2C and S4A, S4C), consistent with the model of CTCF asymmetrical blocking of cohesin “loop extrusion”.

**Figure 2.**
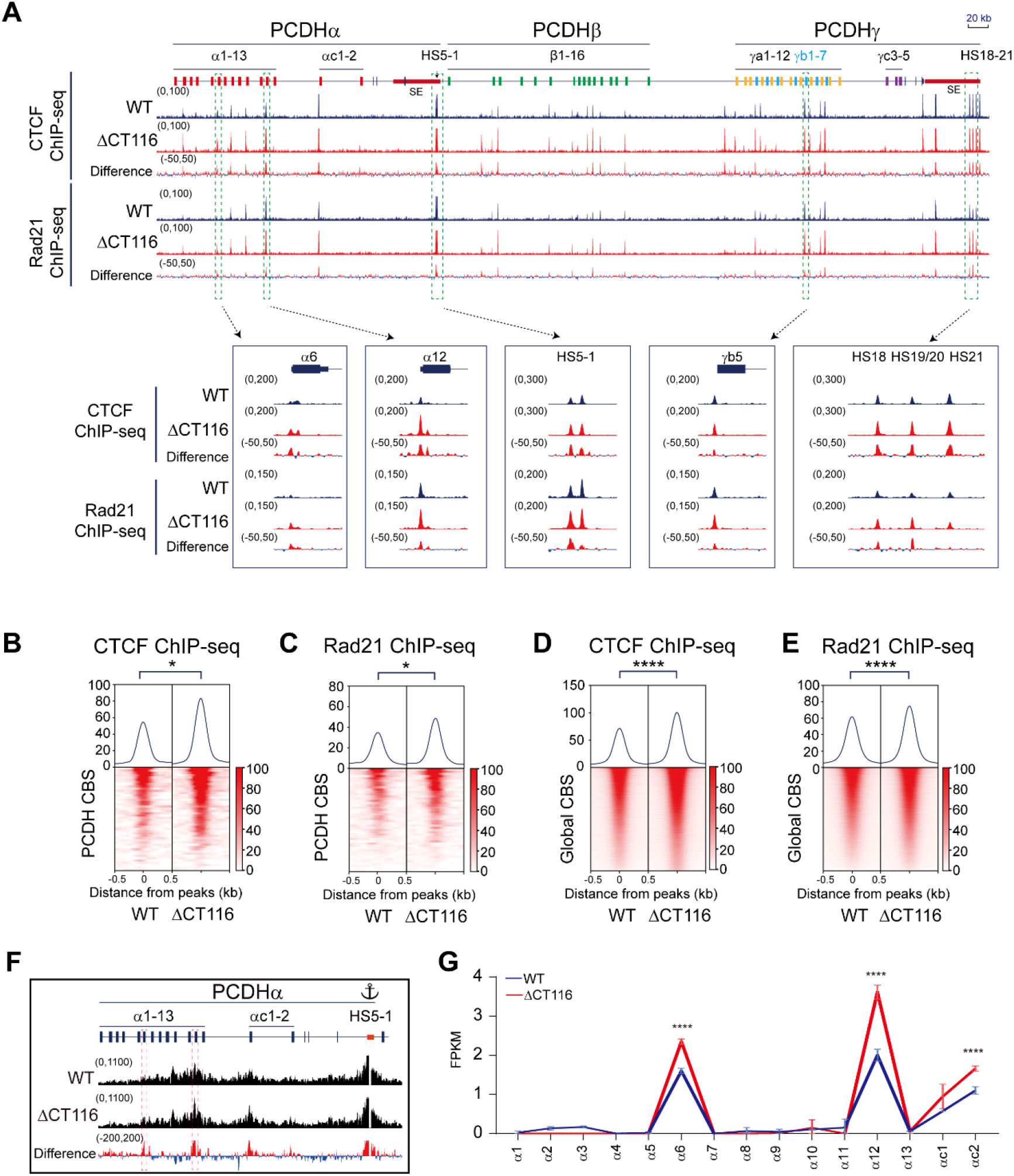
Deleting 116 amino acids from the C-terminus leads to increased CTCF binding and affects gene expression. (A) CTCF and Rad21 ChIP-seq peaks at the *cPCDH* gene complex. (B and C) Heatmaps of CTCF (B) and Rad21 (C) normalized signals at the *cPCDH* CBS elements. Student’s *t* test, \**p* < 0.05. (D and E) Heatmaps of global CTCF (D) and Rad21 (E) ChIP-seq signals. Student’s *t* test, \*\*\*\**p* < 0.0001. (F) QHR-4C interaction profiles of the *PCDHα* gene cluster using *HS5-1* as an anchor. (G) RNA-seq indicates increased expression levels of *PCDHα6, PCDHα12,* and *PCDHαc2.* Data are presented as mean ± SD; Student’s *t* test, \*\*\*\**p* < 0.0001. See also Figures S1, S3, and S4.

We next performed genome-wide analyses and found similar enrichments of CTCF and cohesin upon deletion of the C-terminal 116 AAs (Figure 2D and 2E). Genome-wide analyses of CTCF motifs showed no alteration of all three types of the CBS elements (Figure S4D-S4I). We also performed QHR-4C experiments using the *HS5-1* enhancer as an anchor and found a significant increase of chromatin contacts with the target promoters of *PCDHα6* and *PCDHα12* (Figure 2F). Finally, RNA-seq experiments showed a significant increase of expression levels of *PCDHα6*, *PCDHα12*, and *PCDHαc2* (Figure 2G). These data demonstrated that the unstructured region of C-terminal 116 AAs inhibits CTCF binding to DNA.

### Deletion of the CTCF C-terminal 116 AAs affects gene expression

RNA-seq experiments revealed 432 up-regulated genes (Table S3, Figure S3E) with mean log2FC of 2.36 and 461 down-regulated genes (Table S4, Figure S3F) with mean log2FC of −1.98 in ΔCT116 cell (Figure S3G). Interestingly, we found that the up-regulated genes are closer to increased CTCF peaks (Figure S3H).

### Rescue with series of C-terminal truncated CTCFs reveals negatively-charged region (NCR)

We next generated a series of C-terminal truncated CTCF with V5 tags and transfected them into ΔCTD cells (Figure 3A). Specifically, truncated CTCF with 116 AAs deleted at the C-terminus was constructed as CTCF1-611. In addition, CTCF1-637 contains an additional region of 26 highly-conserved mostly negatively-charged amino acids (NCR). Finally, CTCF1-663 contains a further downstream region of 26 amino acids with 9 proline residues and 7 positively-charged lysine or arginine residues. We generated stable cell lines by infecting with lentiviruses containing these constructs and verified their expression by Western blots (Figure 3A and 3B).

**Figure 3.**
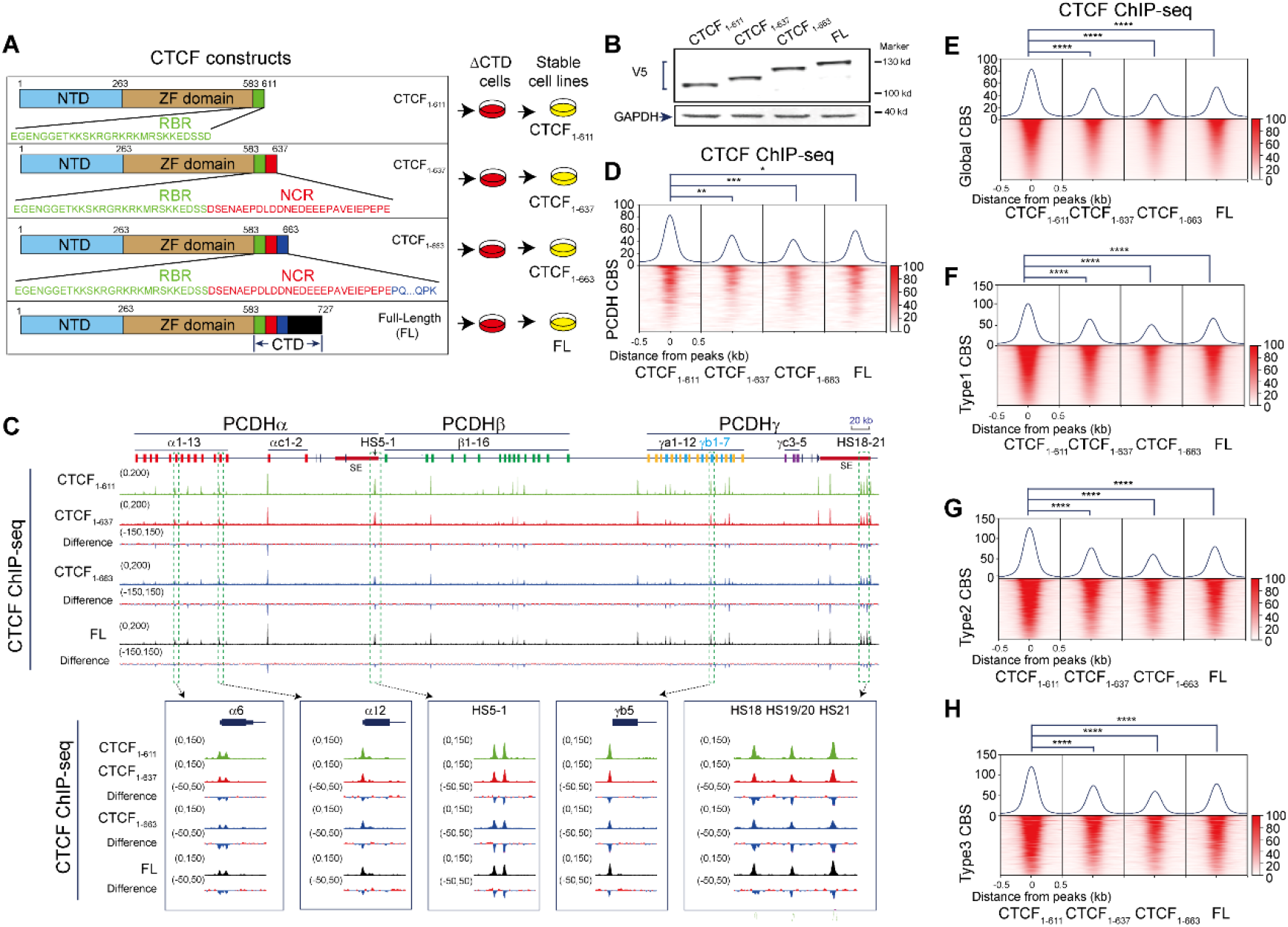
Rescue with a series of C-terminal truncated CTCF proteins reveals a negatively-charged region (NCR) (A) Schematic illustration of different truncated CTCFs used in constructing stable cell lines. All truncated CTCFs are tagged with V5. (B) Western blots of stable cells lines with the V5 antibody indicating truncated CTCF proteins in ΔCTD cells. (C) Binding profiles of different length of CTCFs at the *cPCDH* locus. (D) Heatmaps of CTCF ChIP-seq signals at the *cPCDH* locus. Student’s *t* test, \**p* < 0.05,\*\**p* < 0.01,\*\*\**p* < 0.001. (E) Heatmaps of different truncated CTCFs, showing global binding profiles. Student’s *t* test, \*\*\*\**p* < 0.0001. (F-H) CTCF enrichments at the three types of CTCF motifs. Student’s *t* test, \*\*\*\**p* < 0.0001. See also Figure S5.

We performed ChIP-seq experiments with a specific antibody against V5 tag to investigate CTCF binding profiles and found that the binding strength of CTCF1-611 at the *cPCDH* locus and throughout the entire genome is the highest among all of these four CTCF transgenes including full-length CTCF (Figures 3C-3E and S5A, S5B). Specifically, CTCF1-611 has the highest affinity for all three types of genome-wide CBS elements (Figures 3F-3H and S5C). In conjunction with data of endogenous C-terminal truncation (Figure 2), we concluded that CTCF1-611 has the highest DNA binding affinity and that the negatively charged region (NCR) of 26 AAs from 612 to 637 suppresses CTCF-DNA interactions.

### NCR deletion increases CTCF binding and *cPCDH* expression

To investigate the endogenous function of NCR, we genetically deleted it by screening single-cell CRISPR clones and obtained two cell clones (ΔNCR) (Figure S1H-J). We performed CTCF ChIP-seq experiments with these clones and found that there is a significant increase of CTCF enrichments in the *cPCDH* locus compared with WT controls (Figures 4A, 4B and S6A, S6B). We also performed Rad21 ChIP-seq and found a similar increase of cohesin enrichments at the *cPCDH* locus (Figures 4A, 4C and S6A, S6C). In addition, genome-wide CTCF and cohesin enrichments are also significantly increased upon NCR deletion (Figure 4D and 4E). Furthermore, CTCF or cohesin enrichments are significantly increased at all three types of CBS elements (Figure S6D-S6I). We also performed ChIP-qPCR to further validate increased CTCF binding in ChIP-seq (Figure S6J). We next performed QHR-4C experiments using these single-cell clones with *HS5-1* as an anchor and found there is a significant increase of long-distance chromatin interactions with the target promoters of *PCDHα6* and *PCDHα12* (Figure 4F). Finally, we performed RNA-seq experiments and found that there is a significant increase of expression levels of *PCDHα6* and *PCDHα12* upon NCR deletion. These data suggest an important function of NCR in CTCF binding and gene regulation.

**Figure 4.**
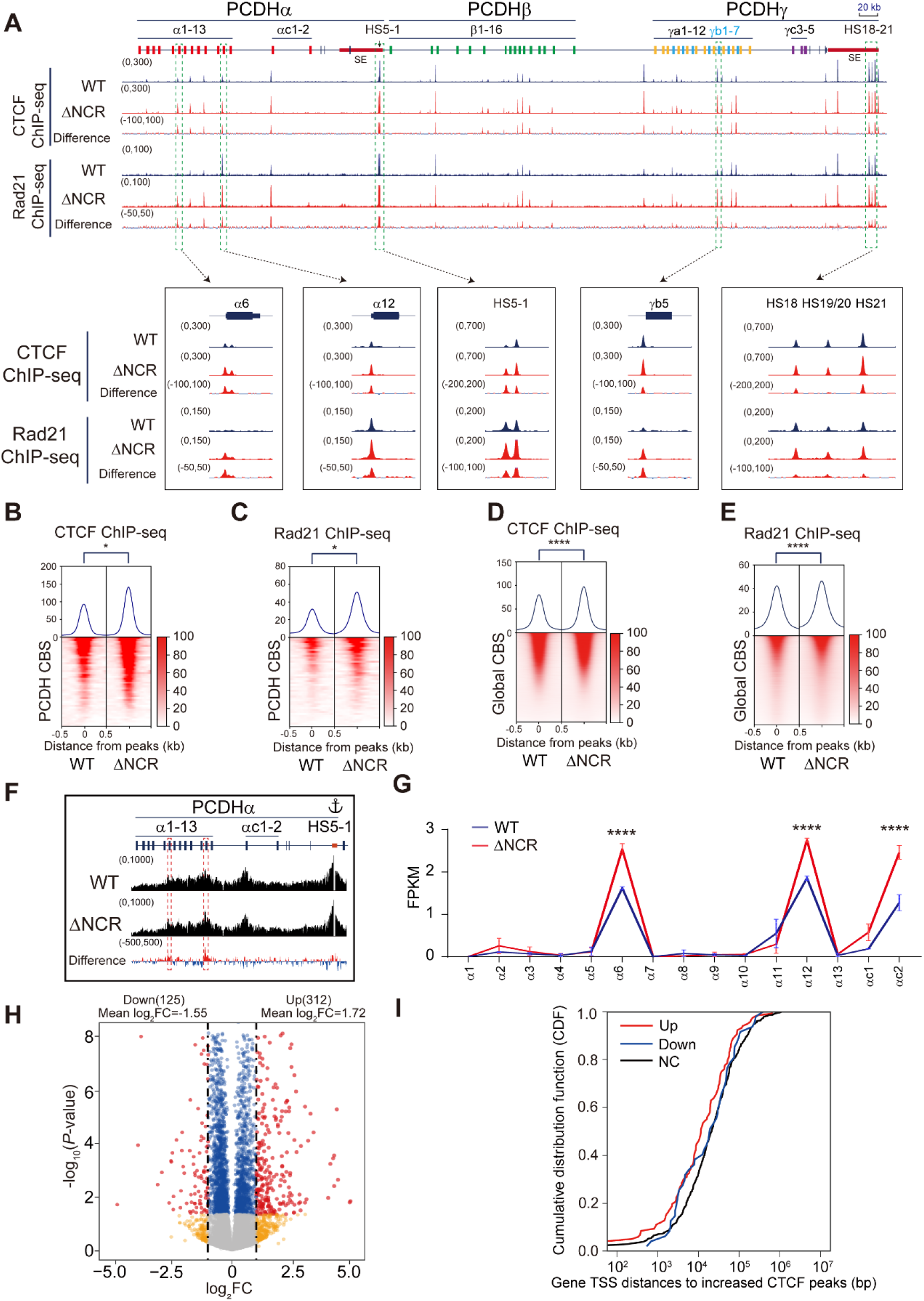
NCR deletion increases CTCF binding and affects gene regulation. (A) CTCF and Rad21 ChIP-seq signals at the *cPCDH* locus, showing increased CTCF and cohesin enrichments at CBS elements upon NCR deletion (Δ612-637). (B and C) Heatmaps of CTCF (B) and Rad21 (C) ChIP-seq signals at the *cPCDH* locus. Student’s *t* test, \**p* < 0.05. (D and E) Heatmaps of CTCF (D) and Rad21 (E) ChIP-seq signals, indicating genome-wide CTCF and Rad21 enrichments upon NCR deletion. Student’s *t* test, \*\*\*\**p* < 0.0001. (F) Quantitative high-resolution 4C (QHR-4C) experiments with *HS5-1* as an anchor, indicating increased chromatin interactions of the *HS5-1* enhancer with *PCDHα6* or *PCDHα12* promoters. (G) RNA-seq indicates increased expression levels of *PCDHα6, PCDHα12,* and *PCDHαc2* upon NCR deletion. Data are presented as mean ± SD; Student’s *t* test, \*\*\*\**p* < 0.0001. (H) Volcano plot of differential gene expression analyses for WT and ΔNCR cells. Red dots, fold change of gene expression upon NCR deletion (log2FC > 1 and adjusted *P* value < 0.05). Blue dots, genes only passed adjusted *P* value < 0.05. Yellow dots, gene only passed log2FC > 1. FC: fold change. (I) TSS distances of up-, down-, or none-regulated (NC) genes to the closest increased CTCF peaks in ΔNCR cells (data are shown as a cumulative distribution function, CDF). See also Figures S1, S3, S6, Table S5, S6.

### Deletion of NCR affects gene regulation

Genome-wide analyses of RNA-seq data identified 312 up-regulated genes (Table S5 and Figure S3I) with mean log2FC of 1.72 and 125 down-regulated genes (Table S6 and Figure S3J) with mean log2FC of −1.55 (Figure 4H). We also found that the up-regulated genes are closer to increased CTCF peaks (Figure 4I). For example, we found that expression levels of *EHD2* (EH domain containing 2) correlates to the change of CTCF binding at the promoter region (Figure S3K-M).

### A role of NCR in rewiring chromatin accessibility

We next performed the assay for transposase-accessible chromatin with sequencing (ATAC-seq) and found that chromatin accessibilities are increased at most ATAC-seq peaks in the *cPCDH* locus upon CTCF NCR deletion (Figure S7). We also observed significantly increased ATAC-seq signals genome wide at increased CTCF peaks (Figure S8A). We next calculated the log2FC of ATAC-seq peaks and found that the majority of ATAC-seq peaks are also increased and minority peaks are decreased (Figure S8B). Specifically, we identified 7373 increased differential accessibility regions (DARs) and 1311 decreased DARs genome wide (Figure S8C). Interestingly, we found that most increased DARs are located at gene promoters and that many decreased DARs are located in the intergenic regions (Figure S8C).

We noted that ∼3/4 DARs (6403) are overlapped with CBS elements and ∼1/4 DARs (2281) are with no CBS (Figure S8D). Among all DARs overlapped with CBS elements, there are 5431 increased DARs and 972 decreased DARs (Figure S8E). We calculated the distance of non-CBS DARs to nearest CBS elements and found that increased DARs are located closer to CBS than decreased DARs (Figure S8F). Integrated analysis with RNA-seq showed that increased DARs correlate with enhanced levels of gene expression (Figure S8G). Together, these data suggest that CTCF NCR regulates chromatin accessibility and gene expression.

### AlphaFold prediction suggests allosteric autoinhibition of CTCF DNA binding by NCR in CTD

We first validated the inhibitory role of NCR in CTCF DNA binding *in vitro* directly by the electrophoretic mobility shift assay (EMSA) (Figure 5A-E). We then used AlphaFold3 to predict the conformation model of the CTCF zinc finger array and CTD with or without DNA ligands (Figure 5F-I).^53^ Intriguingly, AlphaFold3 modeling suggests that the apostate CTCF appears as an autoinhibition conformation with NCR folding back and interacting with the positively-charged surface zinc finger array via electrostatic contacts (Figure 5F). In particular, a cluster of seven negatively-charged residues within NCR appears as a DNA mimicry and can fold onto the positively-charged DNA binding surface of the CTCF zinc finger array via multiple electrostatic interactions (Figure 5G). In addition, both the negatively-charged residues of NCR and positively-charged residues of ZFs are highly conserved across vertebrates (Figure 5H). Binding to DNA targets induces large conformational changes of CTCF and releases the flexible CTD (Figure 5I). Thus, AlphaFold modeling suggests allosteric autoregulation of CTCF DNA binding via DNA-mimicking NCR within the flexible CTD (Figure 5J).

**Figure 5.**
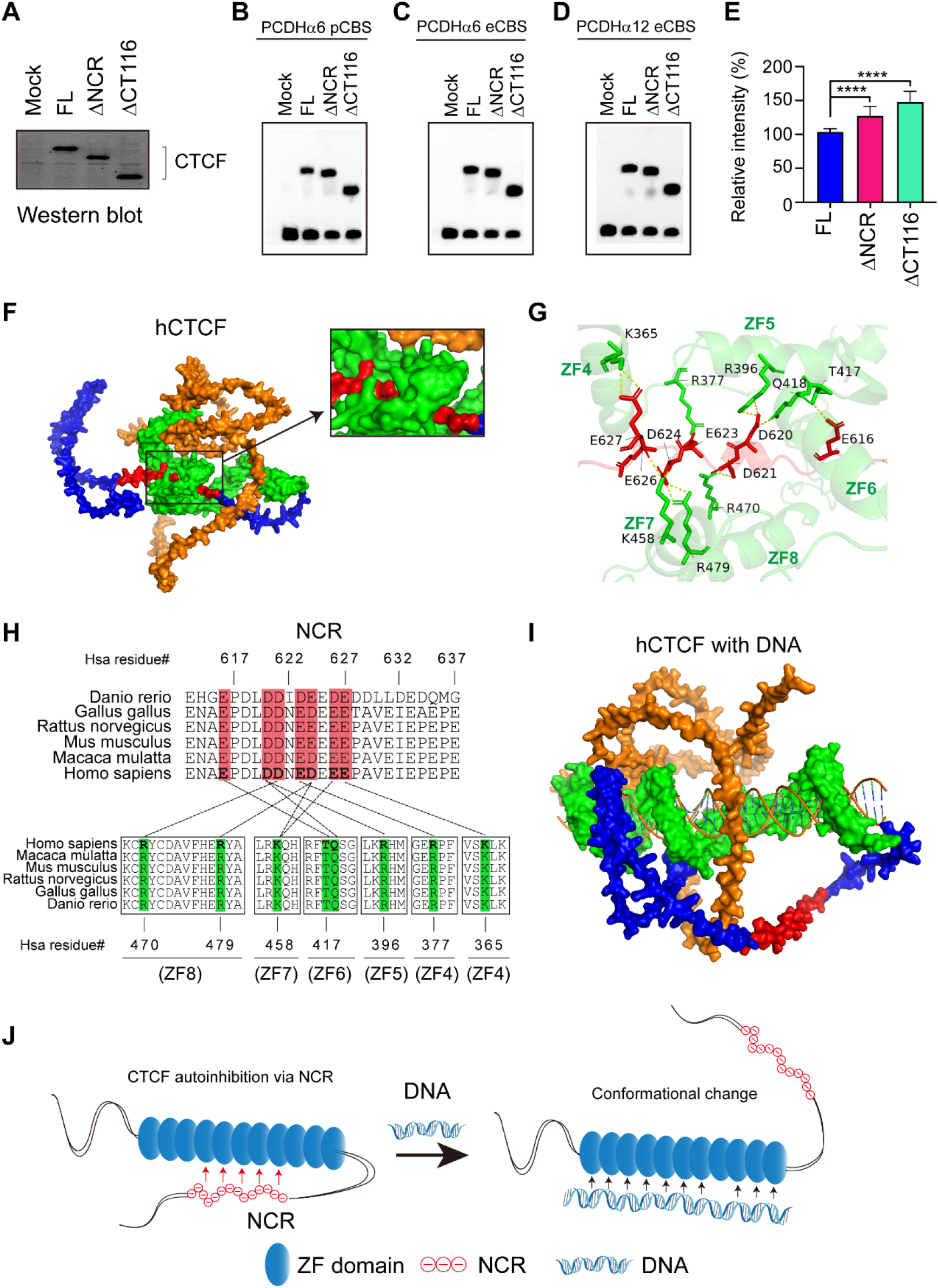
AlphaFold3 modeling suggests autoregulation of CTCF DNA binding by NCR *cis*-interactions. (A) Western blot of recombinant human CTCF proteins. (B-E) EMSA of full-length human CTCF, ΔNCR, ΔCT116 with three different *PCDH* promoter or exonic CBS elements (B-D) with quantifications (E). Data are presented as mean ± SD; Student’s *t* test, \*\*\*\**p* < 0.0001. (F) The surface representation of the AlphaFold3 model of human CTCF in the absence of DNA. NTD, ZF, CTD, and NCR are in orange, green, blue, and red, respectively. (G) Negatively-charged residues of NCR form nine pairs of electrostatic interactions with the zinc finger domain. (H) Pairwise interactions between NCR (top) and ZFs (bottom). (I) The surface representation of the AlphaFold3 model of CTCF with DNA ligand. (J) The model of CTCF autoinhibition via large conformational changes with NCR as a flexible DNA mimicry.

## DISCUSSION

Combinatorial and patterned expression of diverse cadherin-like *cPcdh* genes in single cells in the brain enables a molecular logic of self-avoidance between neurites from the same neurons as well as a functional assembly of synaptic connectivity between neurons of the same developmental origin.^29,54–57^ This complicated *cPcdh* expression program is achieved by ATP-dependent active cohesin “loop extrusion” which brings remote super-enhancer in close contacts to target variable promoters via CTCF-mediated anchoring at tandemly-arrayed directional CBS elements.^14,17–19,27,30,58–60^. In particular, tandem-arrayed CTCF sites function as topological chromatin insulators to balance distance- and context-dependent promoter choice to activate cell-specific gene expression in the brain.^18,19,30,58,59^ Consequently, the dynamic interactions between CTCF and its recognition sites at variable promoters and super-enhancers are central for establishing *cPcdh* expression programs during brain development. In this work we systematically investigated the function of CTCF CTD and uncovered a negatively-charged region (NCR) for maintaining proper CTCF binding to DNA in the large *cPCDH* gene complex and throughout the entire genome.

CTCF is a key 3D genome architectural protein that recognizes a large range of genomic sites via the central domain of 11 tandem zinc fingers and anchors loop-extruding cohesin via the YDF motif of the NTD.^4,12,43–46^ In addition, other zinc-finger architectural proteins such as ZNF143, MAZ, PATZ1, and ZNF263 may collaborate with CTCF to maintain proper CTCF insulation at TAD boundaries.^22,60,61^ The CTCF last ZF and CTD contain an internal RNA-binding region (RBR) of 38 amino acids and this region helps CTCF clustering and searching for authentic CBS elements.^42,52,62,63^ Through a series genetic deletion and rescue experiments, we uncovered an important negatively-charged region (NCR) immediately downstream of the positively-charged RNA-binding region.

Specifically, we showed that NCR deletion leads to a significant increase of CTCF enrichments at all three types of CBS elements throughout the whole genome. In particular, NCR deletion results in increased CTCF binding at *cPCDH* variable promoters and super-enhancers, accompanied by increased chromatin accessibility and long-distance DNA looping. NCR may play an inhibitory role in CTCF clustering via RNA and in dynamic recognition of cognate genomic target sites.^63–65^ NCR either repulses DNA directly or weakens the strength of CTCF interaction with RNA, thus inhibits CTCF clustering and searching for cognate CBS elements. Either way NCR appears to be important in maintaining proper CTCF affinity for cognate genomic sites and specific chromatin looping at the *cPcdh* gene complex and likely throughout the entire genome. Thus, CTCF has an intricate self-adjusting mechanism to control the dynamic binding to genomic sites.

One intriguing allosteric self-adjusting mechanism of CTCF DNA binding is suggested by AlphaFold3 prediction (Figure 5J). According to the large conformational change of CTCF induced by DNA binding, NCR could be released from interacting with the ZF array upon CTCF recognition of genomic CBS elements. Thus, NCR functions as a DNA mimicry and self-associates with the positively-charged surface of the ZF array via electrostatic interactions in the CTCF apostate. Indeed, CTCF CTD interacts with its ZF array in *in-vitro* pull-down experiments.^66^. The self-association of disordered flexible NCR may be a potential general mechanism of DNA and RNA binding proteins. For example, a negatively-charged region of ZNF410 regulates its ZF array to bind DNA via a *cis*-allosteric inhibitory mechanism.^65^ In addition, autoinhibitory intramolecular interactions proof-read U2AF recognition of authentic polypyrimidine tracts during RNA binding.^67^

NCR contains an acidic array of 10 glutamates and 5 aspartates for a total of 15 negatively-charged amino acid residues (Figure 3A). In addition, there are four serine residues immediately upstream of NCR that may be phosphorylated by casein kinase II and thus switching to negative charges upon phosphorylation.^68^ Numerous CTCF mutations are related to multiple cancers or a group of neurodevelopmental diseases known as CTCF-related disorders (CRD).^69,70^ The large spectrum of neurodevelopmental diseases or CTCF-related disorders may be related to dysregulation of clustered protocadherins.^71,72^ Interestingly, mutations within the CTCF NCR which alter its electronic charges are associated with several types of cancers. For example, CTCF mutations of E616K or E626K are associated with melanoma and lung cancers, respectively.^69,70^ The exact pathogenetic mechanisms are not known but are very likely related to disruptions of alternate positive-negative amino acid block patterns of RBR and NCR within CTD and selective partitioning into CTCF-specific trapping zones.^63,73^ Specifically, the positively charged RBR and negatively charged NCR within C-terminal domain of CTCF may constitute recently noticed charge block patterns and may participate in the selective partitioning of phase separation or in the formation of protein condensates during chromatin looping in 3D genome.^73,74^

## Limitations of the study

While our genetic experiments, in conjunction with chromosome conformation capture and RNA-seq, demonstrated that CTCF CTD, in particular NCR, play a crucial role in maintaining proper CTCF binding at the *cPCDH* gene complex, and subsequent *Pcdh* chromatin looping and gene regulation, whether this close correlation between chromatin looping and gene expression could be generalized to the entire genome remains to be tested. In addition, while our ATAC-seq experiments showed that enhanced CTCF DNA binding correlates mostly with higher chromatin accessibility, the exact mechanism is not known but most likely related to pioneering factors.

## RESOURCE AVAILABILITY

### Lead contact

Further information and requests for resources and reagents should be directed to and will be fulfilled by the Lead Contact, Qiang Wu.

### Materials availability

All unique/stable reagents generated in this study are available from the lead contact with a completed materials transfer agreement.

### Data and code availability

- High-throughput sequencing files (ChIP-seq, RNA-seq, ATAC-seq, and QHR-4C) have been deposited into the NCBI Gene Expression Omnibus (GEO) database with the accession number GSE261210, GSE261212, GSE261213, and GSE261209, respectively.
- This paper does not report original code.
- Any additional information required to reanalyze the data reported in this paper is available from the lead contact upon request.

## ACKNOWLEDGEMENTS

We are grateful for advice on bioinformatics from Dr. Jingwei Li and all members of our laboratory for discussion. This study was supported by grants to Q.W. from the National Natural Science Foundation of China (32330016), the National Key R&D Program of China (2022YFC3400200).

## AUTHOR CONTRIBUTIONS

Q.W. conceived the research. Y.Z contributed resources. L.L. performed experiments. L.L. and Y.T. analyzed data. L.L. and Q.W. wrote the manuscript.

## DECLARATION OF INTERESTS

The authors declare no competing interests.

## STAR METHODS

## KEY RESOURCES TABLE

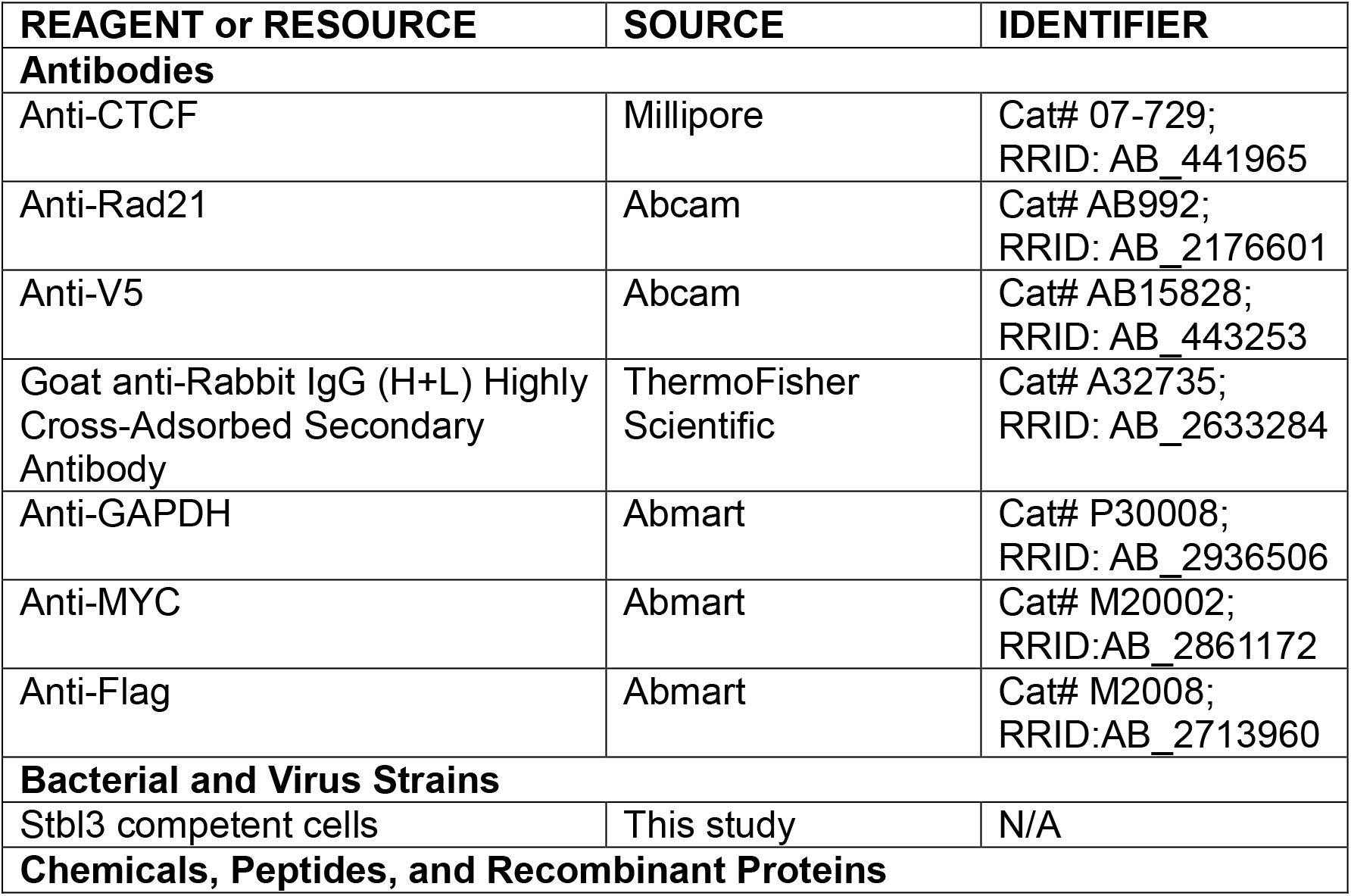
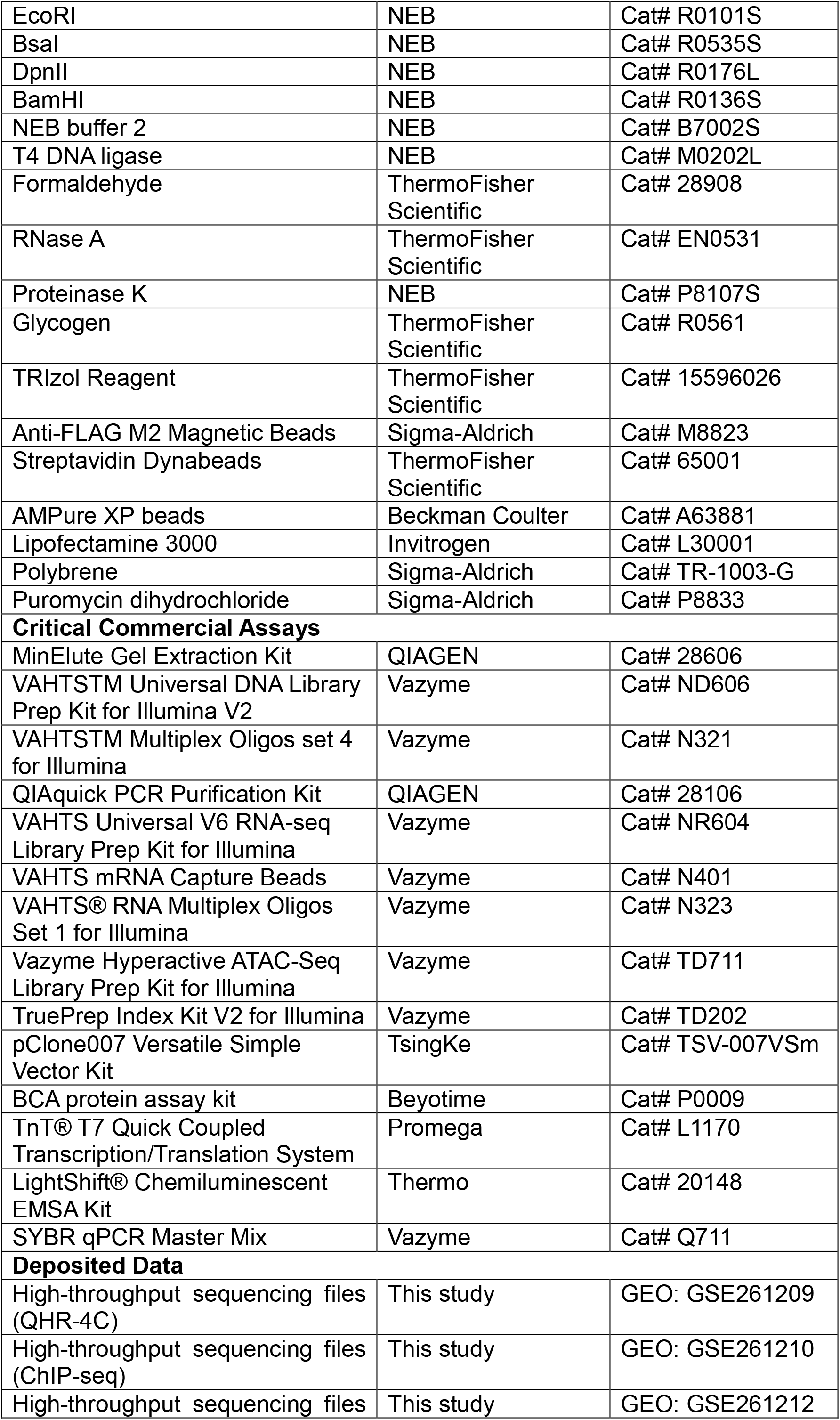
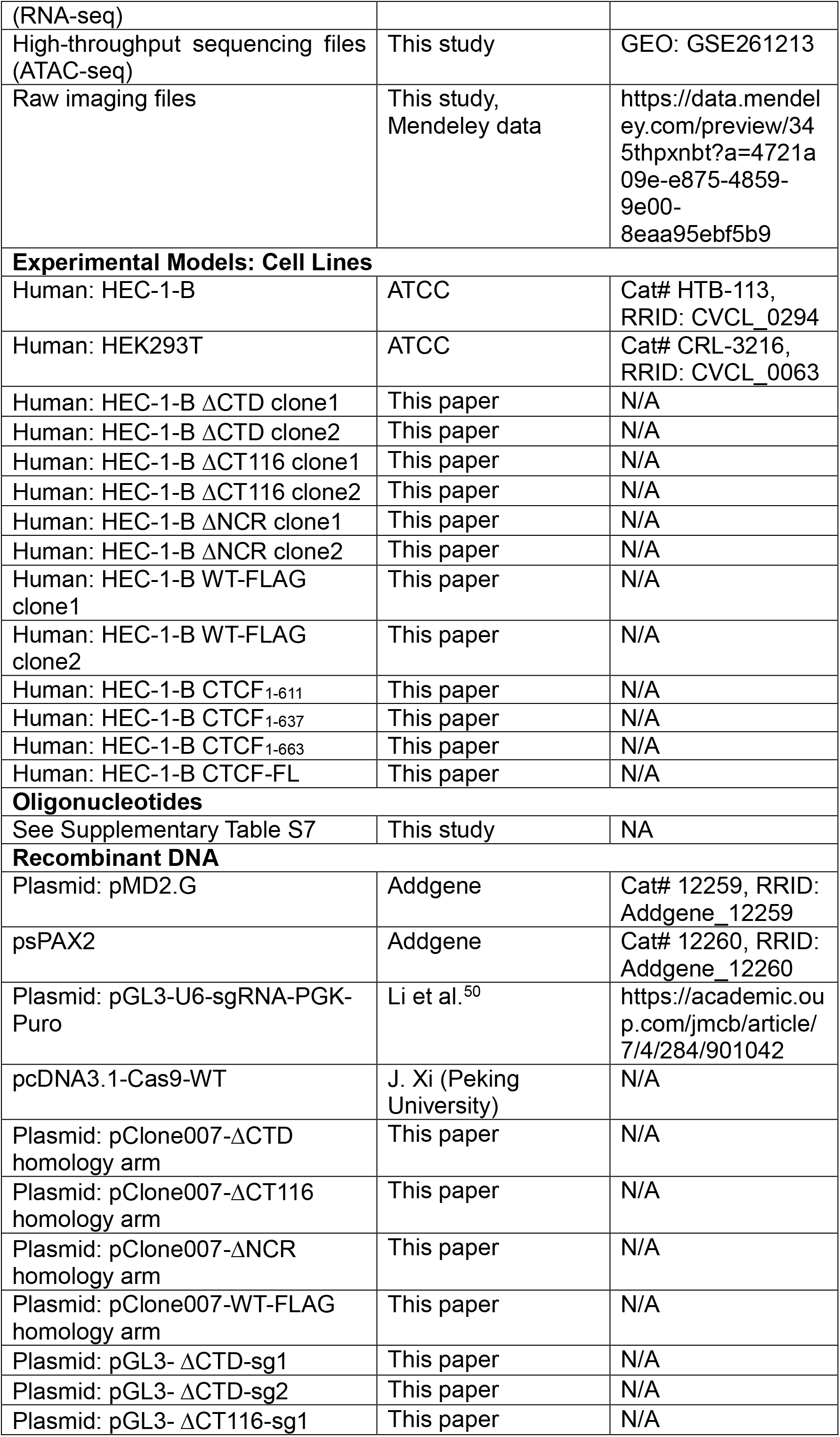
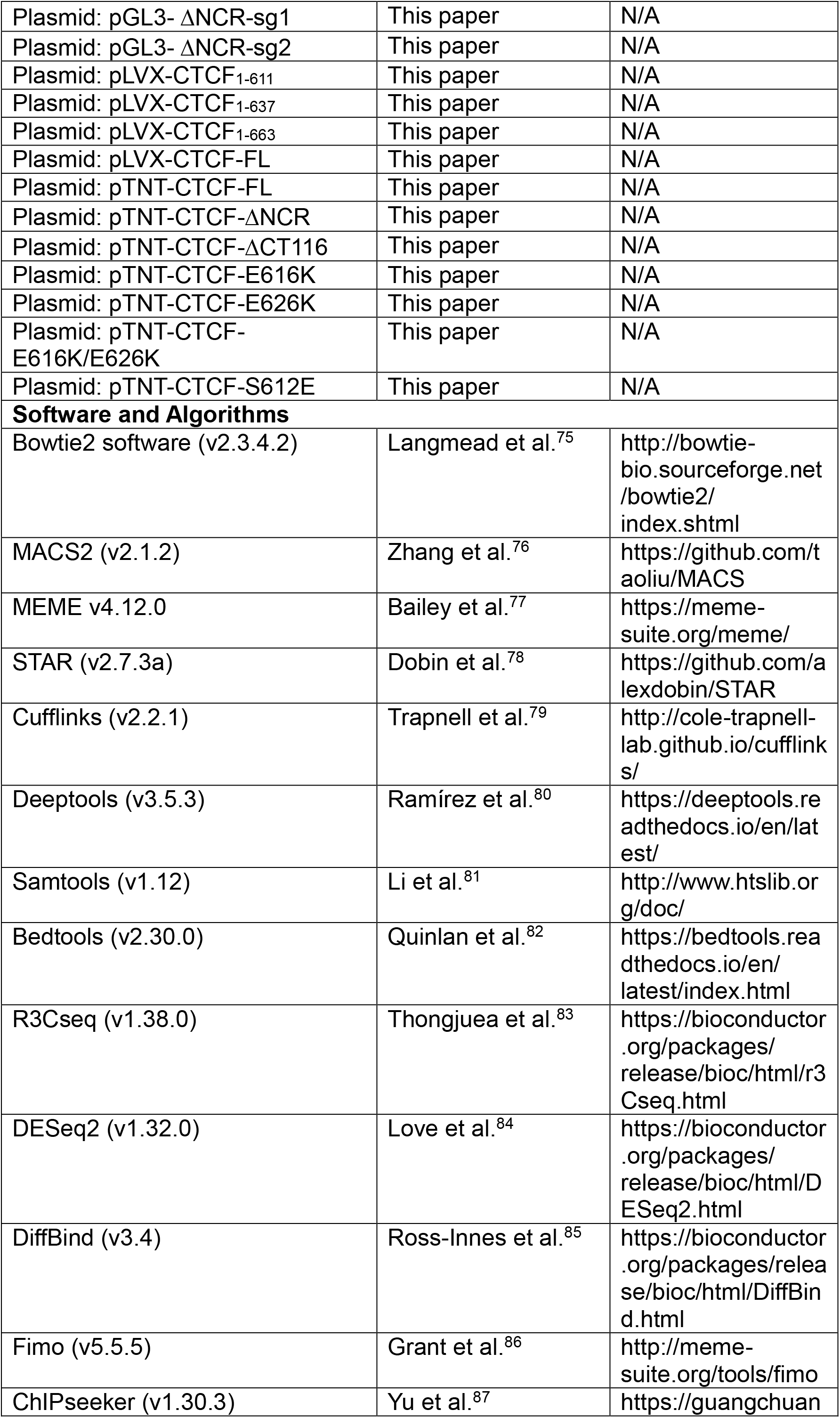
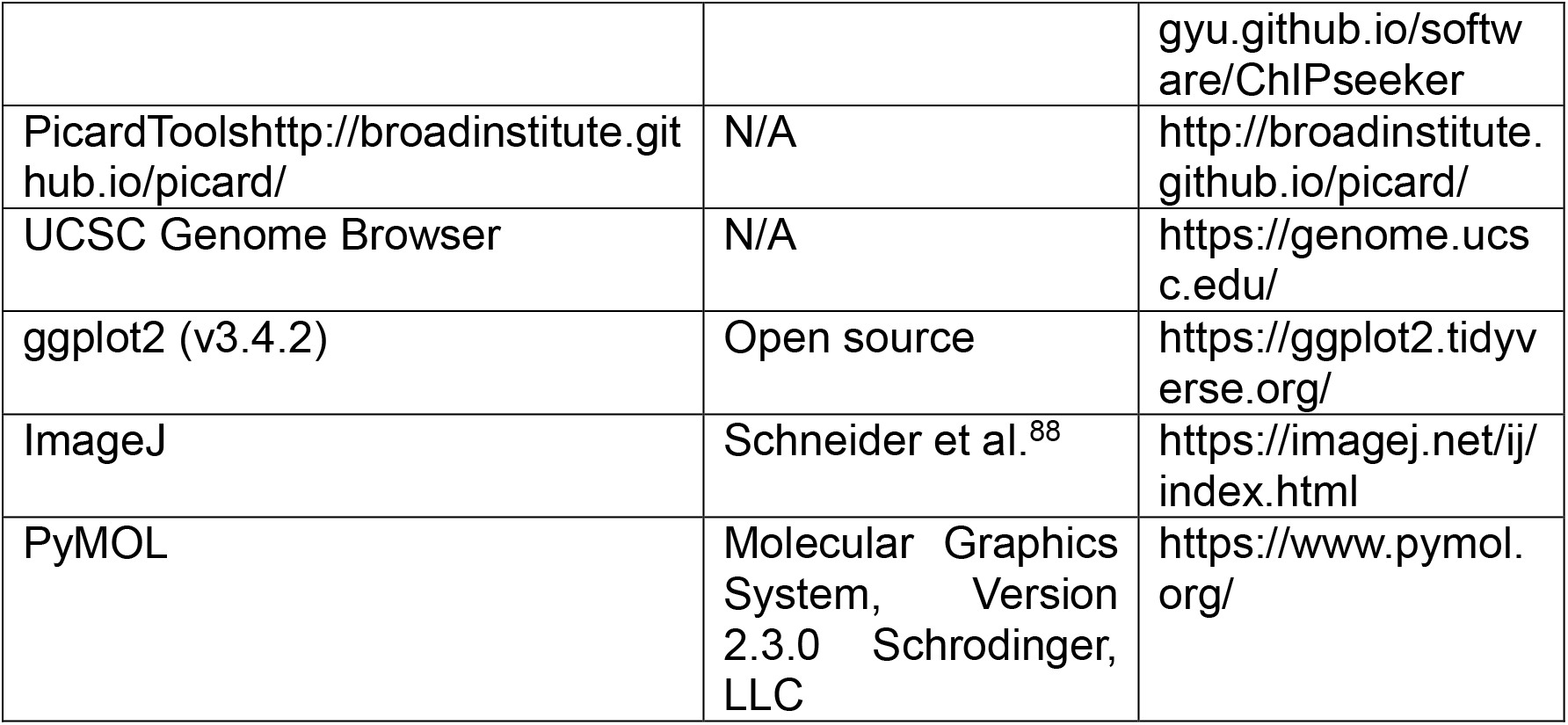

## EXPERIMENTAL MODEL AND STUDY PARTICIPANT DETAILS

### Cells and culture conditions

Human HEC-1-B cells (ATCC) were cultured as previously described in MEM medium (Hyclone) supplemented with 10% (v/v) FBS (Gibco), 2 mM GlutaMAX (Gibco), 1 mM sodium pyruvate (Sigma), and 1% penicillin-streptomycin (Gibco).^17^ Briefly, HEC-1-B cells were maintained at 37 °C in a humidified incubator containing 5% CO2. Medium of cultured cells was changed every 24 h. Cells were passed every 72 h. When cells were passaged, the medium was removed and cells were washed by PBS and then digested by trypsin (Gibco) for 5 min at 37 °C in a humidified incubator containing 5% CO2 and quenched by 10% FBS supplemented PBS. Digested cells were collected by centrifuging at 500 rpm for 5 min at room temperature. Pelleted cells were resuspended with media and seeded to new plates.

HEK293T cells (ATCC) were cultured in Dulbecco’s modified Eagle’s medium (Hyclone) supplemented with 10% (v/v) FBS (Gibco) and 1% penicillin-streptomycin (Gibco). Cells were maintained at 37 °C in a humidified incubator containing 5% CO2. Medium of cultured was changed every 24 h. HEK293T cells were passed every 48 h. The passage process of HEK293T is similar to HEC-1-B except with shortened trypsin digesting time of 2 min.

## METHOD DETAILS

### Plasmid construction

For all CRISPR/Cas9 experiments, the sgRNA plasmids were constructed as described before.^50^ Briefly, pGL3-U6 vector was linearized by BsaI (NEB) to generate the cloning backbone with 5’ overhangs of ‘TGGC’ and ‘TTTG’ at the two ends. In addition, a pair of complementary oligonucleotides (Table S7) containing the sgRNA targeting sequences with 5’ overhangs of ‘ACCG’ or ‘AAAC’ was annealed and ligated to BsaI-digested linearized vector backbone using T4 DNA ligase (NEB). The complete sgRNA sequences are under the control of the U6 promoter and will be transcribed by Pol III in mammalian cells. Finally, the Cas9 plasmid was obtained as a gift from Peking University.

For donor plasmids used in establishing ΔCTD and ΔCT116 stable cell lines via homologous recombination (HR), the donor was designed such that ΔCTD or ΔCT116 were tagged with FLAG sequences (Table S7) for tracing CTCF proteins with FLAG-specific antibodies because deletions of endogenous CTCF C-terminal fragment removed the epitope (659-675 AAs) for CTCF antibodies (Millipore). In addition, we FLAG-tagged WT CTCF via HR at its C-terminus at the endogenous locus as a control. To generate donor plasmid for CRISPR screening single-cell clones through the homologous recombination (HR) pathway, two homologous arms each of ∼1kb flanking the target sites were amplified from genomic DNA by PCR with primers. The amplified homologous arms were PCR-purified and a donor DNA fragment for HR was generated through overlapping-PCR. Finally, the donor DNA fragment was ligated to a T-vector through TA cloning. For ΔNCR cells, the epitope of CTCF antibodies was intact and the donor plasmid was constructed without FLAG.

All rescue CTCF constructs were V5-tagged to distinguish with the endogenous FLAG-tagged CTCF in ΔCTD cells. To construct Lentivirus pLVX vectors containing a series of truncated CTCF C-terminal domains, pLVX vector was first linearized by EcoR1 and BamH1 (NEB) and purified as the cloning backbone. CTCF1-611, CTCF1-637, CTCF1-663, and full length CTCF were cloned from a cDNA library of HEC-1-B cells with the same 5’ primers containing EcoRI restriction endonuclease sites and different 3’ primers containing BamHI restriction endonuclease sites in conjunction with the V5 tag sequence (Table S7). Truncated CTCF fragments amplified from cDNA were digested with restriction endonucleases and ligated into pLVX linearized vector using T4 DNA ligase.

### ChIP-seq

Chromatin immunoprecipitation followed by high-throughput sequencing (ChIP-seq) experiments were performed as described before.^14^ Briefly, 5×10^6^ cells were collected and crosslinked with 1% formaldehyde, quenched by 2 M glycine at a final concentration of 125 mM and washed by ice-cold PBS twice. Crosslinked cells were lysed twice by the ChIP buffer (10 mM Tris-HCl pH 7.5, 1 mM EDTA, 1% Triton X-100, 0.1% SDS, 0.1% sodium deoxycholate, 0.15 M NaCl, and 1×protease inhibitors, Roche) with slow rotation at 4 °C for 10 min. The lysed cells were centrifuged at 2,500 g at 4 °C for 10 min to isolate cell nuclei. The isolated nuclei were resuspended with the ChIP buffer and sonicated using a Bioruptor Sonicator (with the high energy setting at a train of 30-s sonication with 30-s interval for 30 cycles) to fragment DNA to 200-400 bp. The sonicated mixture was centrifuged at 14,000 g at 4 °C for 30 min and the supernatant was then precleared by protein A/G magnetic beads (Thermo 26162) for 3h at 4 °C with slow rotation. Antibody (CTCF: Millipore 07-729, Rad21: Abcam ab992, V5: Abcam ab15828) or Anti-FLAG antibody conjugated Magnetic Beads (Sigma M8823) were added to the precleared solution and incubated at 4 °C overnight with slow rotation to precipitate CTCF or cohesin protein-DNA complex.

Protein A/G beads were added and incubated for 3 h at 4 °C with slow rotation to capture the antibody-protein-DNA complex. The ChIP buffer, high salt buffer (10 mM Tris-HCl pH 7.5, 1 mM EDTA, 1% Triton X-100, 0.1% SDS, 0.1% sodium deoxycholate, 0.4 M NaCl), no salt buffer (high salt buffer without NaCl), LiCl buffer (50 mM HEPES pH 7.5, 1 mM EDTA, 1% NP-40, 0.7% sodium deoxycholate, 0.5 M LiCl), and 10 mM Tris-HCl buffer (pH 7.5) were used sequentially to wash the beads. Finally, the elution buffer (50 mM Tris-HCl pH 8.0, 10 mM EDTA, 1% SDS) was used to elute ChIP DNA from beads at 65 °C for 1h with 1,000 rpm shaking. The eluted protein-DNA complex was reverse-crosslinked at 65 °C overnight with 1,000 rpm shaking to dissociate DNA. Finally, the proteinase K (NEB) was added and incubated for 2 h at 55 °C to digest protein and the RNase A (Thermo) was added and incubated for 2h at 37 °C to digest RNA.

The DNA was purified by adding equal volume of phenol-chloroform and mixed by vigorously shaking. The mixture was centrifuged at 4 °C at 14,000 g for 10 min to separate the proteins and DNA. The supernatant containing DNA was transferred to a new tube. 2.5-fold volume of ice-cold ethanol, 1/10 volume of 3 M NaAc (pH 5.2), and 1.5 μl of glycogen (Thermo) were added to participate DNA at −80 °C for 1 h. The sample was centrifuged at 14,000 g at 4 °C for 30 min to pellet DNA. 70% ethanol was added to wash DNA pellets and finally the sample was centrifuged at 14,000g at 4 °C for 10 min. The supernatant was removed and the pelleted DNA was air-dried for 5 min to remove residual ethanol. The DNA was then dissolved in the nuclease-free water. The concentration of DNA was measured using Qubit (Invitrogen).

To prepare DNA library for deep sequencing, we used the Universal DNA Library Prep Kit (Vazyme ND606). Briefly, DNA was first end-repaired and ligated to adapters from the Multiplex Oligos Set (Vazyme, N321). Adapter-ligated DNA was then purified using AMPure XP beads (Beckman) and the final ChIP library was amplified by PCR. The library was sequenced on an Illumina NovaSeq platform. All of the ChIP-seq experiments were performed with at least two biological replicates.

### ChIP-qPCR

The chromatin immunoprecipitation followed by quantitative PCR (ChIP-qPCR) experiments were performed as described before.^14^ ChIP steps were the same as the ChIP-seq experiments described above. PCR primers were designed around CTCF binding sites (Table S7). qPCR were then carried out on the ABI QS6 platform using the SYBR qPCR master mix.

### Western blot

1×10^6^ cells were collected and lysed by the RIPA buffer (50 mM Tris-HCl pH 7.4, 150 mM NaCl, 1% Triton X-100, 1% sodium deoxycholate, 0.1% SDS, and 1× protease inhibitors) on ice for 30 min. The sample was sonicated and centrifuged at 14,000 g for 15 min at 4 °C to remove cell debris. The concentration of supernatant proteins mixture was measured using the BCA protein assay kit (Beyotime). The supernatant proteins were mixed with 5× SDS protein loading buffer with DTT and incubated at 95 °C for 10 min to denature the proteins. The denatured protein mix was centrifuged at 14,000g for 15 min at room temperature and separated on SDS-PAGE.

The separated proteins in gel were transferred to the nitrocellulose membrane through electrophoreses. After transferring, the membrane was blocked by 5% non-fat milk in PBST for 2 h at room temperature and then washed 3 times with PBST. Corresponding antibodies were incubated with the membrane overnight at 4°C with slow shaking. The membrane was washed 3 times with PBST. The secondary antibody (Invitrogen) was then incubated with the membrane for 90 min at room temperature with slow shaking. Finally, the membrane was washed by PBST for 3 times and scanned by the Odyssey System (LI-COR Biosciences). The intensity of bands were measured by ImageJ.

### Recombinant CTCF Protein Production

The recombinant mutation proteins of CTCF were prepared by rabbit reticulocyte lysate (Promega L1170) as previously described.^17^ Firstly, we linearized the empty pTNT plasmid by EcoR1 and Not1 (NEB). Then we cloned different CTCF deletions of the CTCF C-terminal domain from cDNA through PCR. The PCR products containing EcoR1 and Not1 restriction endonuclease sites were gel-purified and digested by EcoR1 and Not1. The digested DNAs were ligated to linearized pTNT vectors using T4 DNA ligase. The plasmids were used as a template to express CTCF mutants using the rabbit reticulocyte lysate at 30°C for 90 min. The recombinant CTCF proteins were finally analyzed by Western Blots.

### Electrophoretic Mobility Shift Assay (EMSA)

The EMSA experiments were performed as described before.^17^ We first cloned the probe sequences containing CBS from genomic DNA. Then the DNAs were gel-purified and ligated into a T-vector. The sequences were verified by Sanger sequencing. We finally preformed PCR experiments using plasmids as a template with 5’ biotin labeled primers. We used high-fidelity DNA polymerase to perform PCR and PCR products were gel-purified as EMSA probes. The concentrations of probes were measured by Nano-drop. We performed the EMSA experiments using LightShift Chemiluminescent EMSA reagents (Thermo) according to the manuals. Briefly, equal amounts of protein were incubated in the binding buffer (containing 10 mM Tris, 50 mM KCl, 5 mM MgCl2, 0.1 mM ZnSO4, 1 mM dithiothreitol, 0.1% (v/v) Nonidet P-40 (NP-40), 50 ng/µl poly (dI-dC), and 2.5% (v/v) glycerol) on ice for 20 min to reduce the background. We added the same amounts of probes to the binding buffer and then incubated at room temperature for 30 min. The binding mixtures were then electrophoresed on 5% non-denaturing polyacrylamide gels in the ice-cold 0.5×TBE buffer (45 mM Tris-borate, 1 mM EDTA, pH8.0), and then the gels were transferred to nylon membranes. The membranes were cross-linked under UV for 12 min. We incubated the membrane in the blocking buffer for 20 min. The membranes were treated with streptavidin-horseradish peroxidase conjugate for 20 mins. We washed the membranes for 4 times using the wash buffer and stained the membrane by chemiluminescence using the ChemiDoc XRS+ system (Bio-Rad). The intensity of bands was measured by the ImageJ software. All of the EMSA experiments were performed with at least two biological replicates.

### CRISPR genome editing and single-cell cloning

CRISPR genome editing was performed as described.^50,51^ Cas9 and sgRNA plasmids, and donor constructs were transfected in 12-well plates at 70 % confluency using lipofectamine 3000 (Invitrogen). Medium containing lipofectamine 3000 and plasmid DNA was then replaced with fresh medium after 6 h. After transfection for 48 h, the transfected cells were incubated with medium containing 2 μg/ml puromycin and refreshed daily. After 4 days, the cells were recovered for 2 days in fresh medium without puromycin. The cells were then dissociated by trypsin and seeded to 96-wells cell plates at the concentration of about one cell per well. After incubation for about 1 week in 96-well plates, the seeded cells were checked under microscope and single-cell clones were marked manually.

PCR genotyping was performed when seeded single-cell clones reached ∼80% confluence in 96-well plates. The cells were digested using trypsin and PCR-genotyped with a pair of primers matching sequences outside of the homologous arms (Table S7). The positive PCR products were purified and sequenced by Sanger sequencing for confirmation. The identified single-cell clones were incubated for another 4∼5 passages and confirmed again by PCR genotyping and Sanger sequencing.

Given that the CTCF antibody used in ChIP-seq recognizes 659-675 AAs of CTCF, we added a FLAG tag in ΔCTD and ΔCT116 cell lines via homologous recombination. We also inserted a FLAG tag in WT cell as an experimental control. For ΔCTD cells, we genotyped 191 single-cell clones and found 3 single-cell clones with CTD deletion. For WT-FLAG cells, we genotyped 160 single-cell clones and found 4 of them with FLAG tag insertion. For ΔCT116 cells, we genotyped 192 single-cell clones and found 8 of them with CT116 deletion. For ΔNCR cells, we genotyped 240 single-cell clones and found 6 of them with NCR deletion. For the cells with same genotype, we used two independent single-cell clones for subsequent experiments.

### Lentivirus packaging

Lentiviruses were used to build stable transgenic cell lines that expressing different CTCF rescue constructs in ΔCTD cells. pLVX plasmids containing truncated CTCF sequences and puromycin selectable marker, and helper plasmids (psPAX2, pMD2.G, Addgene) were transfected into HEK293T cells in 6-well plate at 70% confluence using lipofectamine 3000 reagents. The medium containing lipofectamine and plasmid DNA was replaced with fresh medium after 6 h. The medium containing lentivirus was harvested after transfection for 48 h for the first time, and then replaced with fresh medium and harvested again after another 24 h. The harvested medium was then centrifuged at 1,300 rpm at 4 °C for 5 min to remove cells and then filtered by 0.45 μm filter (Merck) and the virus was collected and stored at -80 °C.

### Lentivirus infection and stable cell line establishment

All lentivirus particles expressing different truncated CTCF constructs were thawed on ice and infected with ΔCTD cells by adding polybrene at 8 ug/ml. After infection for two days, the infected cells were incubated with medium containing 2 μg/ml puromycin for 4 days and replaced daily. After selection, cells were incubated with fresh medium without puromycin for 2 days for recovering. The cells were then cultured for another 12 days and harvested for Western blot and ChIP-seq experiments.

### RNA-seq

The RNA-seq experiment was performed as previously described.^17^ Briefly, about 1×10^6^ cells in 6-well plates were collected and washed by ice-cold PBS for 3 times. One ml of TRIzol (Invitrogen) was added to the cells and mixed thoroughly. After incubation for 5 min at room temperature, 0.2 ml of chloroform was added and mixed well by hands. The sample was then incubated for 2 min at room temperature and centrifuged at 12,000 rpm for 10 min at 4 °C. The supernatant was transferred to a new tube and mixed with equal volume of isopropanol. The sample was incubated at room temperature for 10 min and centrifuged at 12,000 rpm for 15 min at 4 °C to pellet RNA. After removing the supernatant, 1 ml of 75% ethanol was added to wash RNA. Finally, RNA pellets were dissolved in nuclease-free water. The quantity and concentration of the total RNA was measured by NanoDrop (Thermo).

The Oligo (dT) coupled magnetic beads (Vazyme) were used to purify mRNA from total RNA. The mRNA library preparation was performed using the Universal V6 RNA-seq Library Prep Kit (Vazyme) according to the manual. Briefly, mRNA was fragmented by heating at 94 °C for 8 min. Next, reverse transcription for the first-strand cDNA was performed and then the double-strand cDNA was synthesized. Adapters from the RNA Multiplex Oligos Set (Vazyme) were added to cDNA and the adapter-ligated cDNA was purified by AMPure XP beads (Beckman). Finally, Library was amplified through PCR and was sequenced on an Illumina NovaSeq platform. All of the RNA-seq experiments were performed with at least two biological replicates.

### ATAC-seq

ATAC-seq was performed as previously described.^59^ Briefly, cells in 12-well plates at 80% confluence were digested by trypsin and were collected by centrifuging at 500 rpm at room temperature for 5 min. 50,000 cells were collected and washed twice by PBS. The cells were then washed twice by ice-cold buffer by centrifuging at 2,300 rpm at 4 °C for 5 min. The cells were then lysed by fresh prepared lysis buffer containing NP-40, Tween-20, and digitonin on ice for 5 min and centrifuged for 10 min at 2,300 rpm at 4 °C to collect the cell nuclei.

The chromatins were fragmented with Tn5 transposase and sequence adaptor at 37 °C for 30 min. The fragmentation reaction was terminated by adding the stop buffer at room temperature for 5 min. The fragmented DNA was purified by ATAC DNA Extract Beads (Vazyme). Finally, PCR was performed to amplify the library using primers in TruePrep Index Kit V2 (Vazyme) and the library was sequenced on an Illumina NovaSeq platform. All of the ATAC-seq experiments were performed with at least two biological replicates.

### QHR-4C

Quantitative high-resolution circularized chromosome conformation capture (QHR-4C) experiments were performed as previously described.^19^ Briefly, 1×10^6^ cell were collected and crosslinked by 2% formaldehyde at room temperature for 10 min with slow rotation. The crosslinking reaction was quenched by adding 2 M glycine to the final concentration of 200 mM. The crosslinked cells were washed twice by ice-cold PBS by centrifuging for 5 min at 800 g at 4 °C and then lysed twice by ice-cold lysis buffer (50 mM Tris-HCl pH 7.5, 150 mM NaCl, 5 mM EDTA, 0.5% NP-40,1% Triton X-100, and 1× protease inhibitors) for 10 min at 4 °C with slow rotation. The cell nuclei were collected by centrifuge at 800 g at 4 °C for 5 min. The pelleted nuclei were resuspended in 73 μl of nuclease-free water, 10μl of DpnII buffer (NEB), and 2.5 μl of 10% SDS and incubated for 1 h at 37 °C with shaking at 900 rpm. 12.5 μl of 20% Triton X-100 was added for 1 h at 37 °C with shaking at 900 rpm. 2 μl of DpnII (NEB) was added to digest the chromatin overnight at 37 °C with shaking at 900 rpm and deactivated at 65 °C for 20 min.

The nuclei were then collected through centrifuging at 1,000 g for 1 min. The supernatant was carefully removed. The proximity ligation was performed with resuspended nuclei by adding 100μl of T4 ligation buffer (NEB) containing 1 μl of T4 ligase and incubated at 16 °C for 24 h. 1 μl of proteinase K was then added to the ligation mixture to digest the proteins and incubated for 4 h at 65 °C for reverse crosslinking. The DNA was purified by phenol-chloroform as described above in ChIP-seq. Finally, the purified DNA was dissolved in 50 μl nuclease-free water and sonicated with the Bioruptor Sonicator (with the low energy setting by a train of 30-s sonication with 30-s interval for 12 cycles) to fragment DNA to sizes of 200-600 bp.

The PCR amplification was performed using 5’ biotin-labeled primers (Table S7) to capture DNA anchored at the *HS5-1* enhancer. To maximize the PCR product, 100 μl of reaction system and 60 cycles of PCR were used. The PCR product was incubated at 95 °C for 5 min and immediately chilled on ice to obtain single-strand DNA (ssDNA). The biotin-labeled ssDNA was collected by incubating with Streptavidin Beads (Invitrogen) for 2 h at room temperature and the beads were washed twice by the washing buffer (5 mM Tris-HCl pH7.5, 1 M NaCl, 0.5 mM EDTA).

To prepare library for deep sequencing, adapters containing sequences that match the 3’ end of the Illumina P7 sequence were ligated to ssDNA at 16 °C for 24 h. The adapters were generated through annealing of two complementary primers (Table S7) in the annealing buffer (25 mM NaCl, 10 mM Tris-HCl pH 7.5, 0.5 mM EDTA). The beads with ssDNA-adapter were washed twice by the washing buffer. Finally, the library was amplified through PCR with two primers. The forward primer contains the Illumina P5 sequences and the sequences adjacent to the *HS5-1* anchor. The reverse primer contains the Illumina P7 sequences and indexes. The amplified library was sequenced on an Illumina NovaSeq platform. All of the QHR-4C experiments were performed with at least two biological replicates.

### Data analysis of ChIP-seq

Raw FASTQ files were aligned to the human reference genome (GRCh37/hg19) using Bowtie2.^75^ The MarkDuplicates module of PICARD tools (http://broadinstitute.github.io/picard/) was used to remove the duplicates and Samtools^81^ was used to index or sort bam files. ChIP-seq peaks were called by MACS2^76^ with the default parameter. The reads counts were normalized to reads per kilobase per million mapped (RPKM) using bamCoverage module of Deeptools^80^ with a bin sized of 20 bp. The plotHeatmap module of Deeptools was used to generate heatmaps. Normalized read counts were converted to bedGraph to be visualized in the UCSC genome browser. Differential binding analyses were performed using DiffBind with the default parameter.^85^ Motif analyses of CTCF were performed using the MEME suite^77^ and FIMO.^86^

### Data analysis of RNA-seq

Raw FASTQ files were aligned to the human reference genome (GRCh37/hg19) using STAR^78^ with default parameters and the RPKMs were calculated using Cufflink.^79^ Differential analysis of gene expression was performed using DEseq2.^84^ Volcano plot was generated by ggplot2. TSS distance to nearest CTCF peaks were calculated by Bedtools.^82^

### Data analysis of ATAC-seq

Raw FASTQ files were aligned to the human reference genome (GRCh37/hg19) using Bowtie2.^75^ ATAC-seq peaks were called by MACS2.^76^ Differential accessibility regions (DARs) were analyzed using DESeq2.^84^ ATAC-seq peak annotation was performed with ChIPseeker.^87^ Bedtools was used in overlapping analyses of DAR and CBS elements and in calculating the distance of DARs to nearest CBS elements.^82^

### Data analysis of QHR-4C

Raw FASTQ files were aligned to the human reference genome (GRCh37/hg19) using Bowtie2.^75^ Reads were normalized by r3Cseq program (version 1.20).^83^

## QUANTIFUCANTION AND STATISTICAL ANALYSIS

All statistical tests were performed with the *R* scripts. Statistical significance values were calculated using the Student’s *t* test. *p* < 0.05 was shown as *, *p* < 0.01 was shown as **, *p* < 0.001 was shown as *** and *p* < 0.0001 was shown as ****.

---

**Figure S1.**
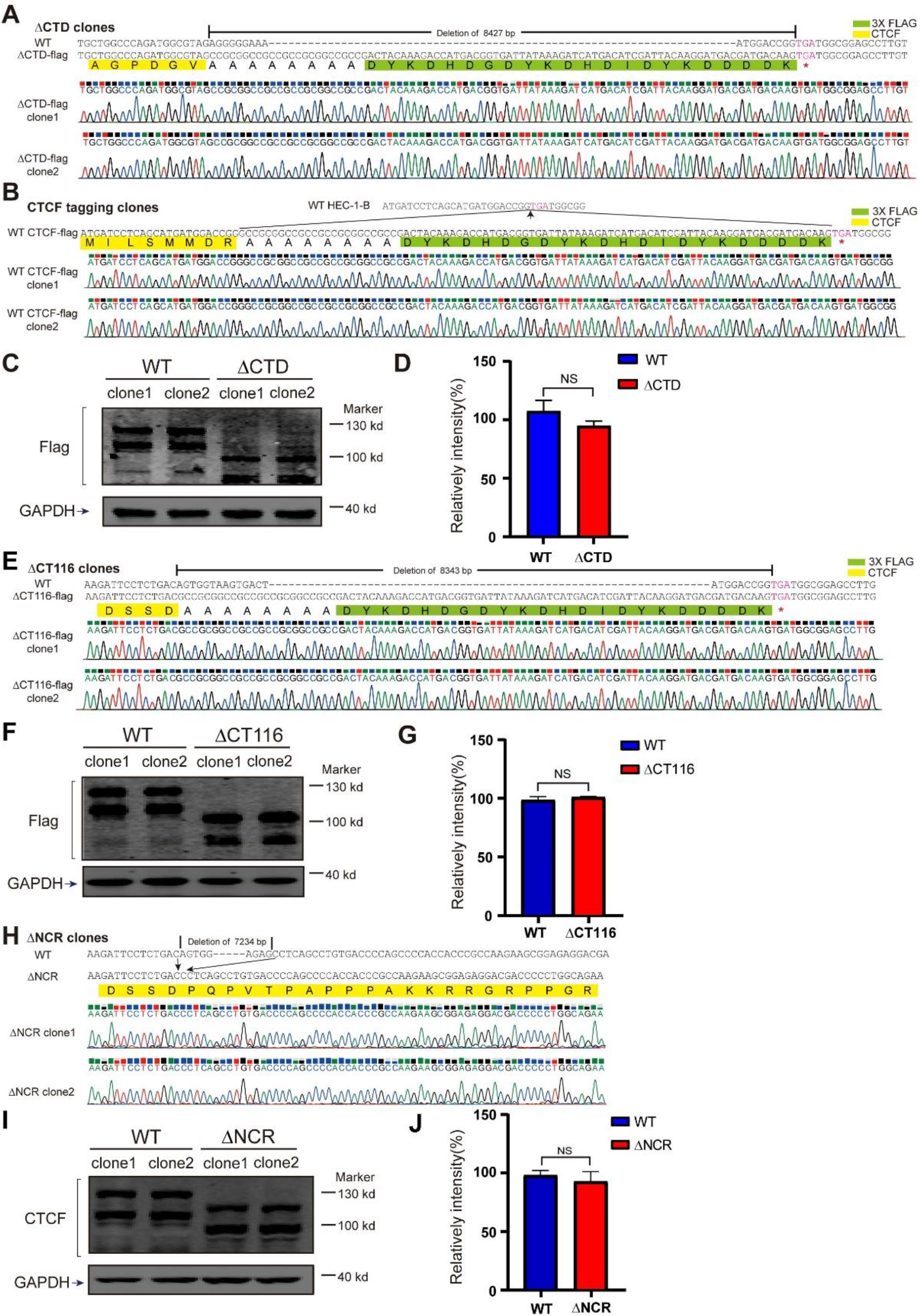
Genotyping and Western blots of single-cell CRISPR clones of CTD, CT116, or NCR deletion as well as wild-type CTCF FLAG-tagging cell clones. Related to Figures 1, 2, and 4. (A-D) Genotyping of the two ΔCTD-FLAG (A), WT-FLAG (B) singe-cell clones by Sanger sequencing, and their Western blots (C) with quantification (D). (E-G) Genotyping of the two ΔCT116-FLAG singe-cell clones (E) by Sanger sequencing, and their Western blots (F) with quantification (G). (H-J) Genotyping of the two ΔNCR singe-cell clones (H) by Sanger sequencing, and their Western blots (I) with quantification (J). Data are presented as mean ± SD. Student’s *t* test, NS, *p* > 0.05.

**Figure S2.**
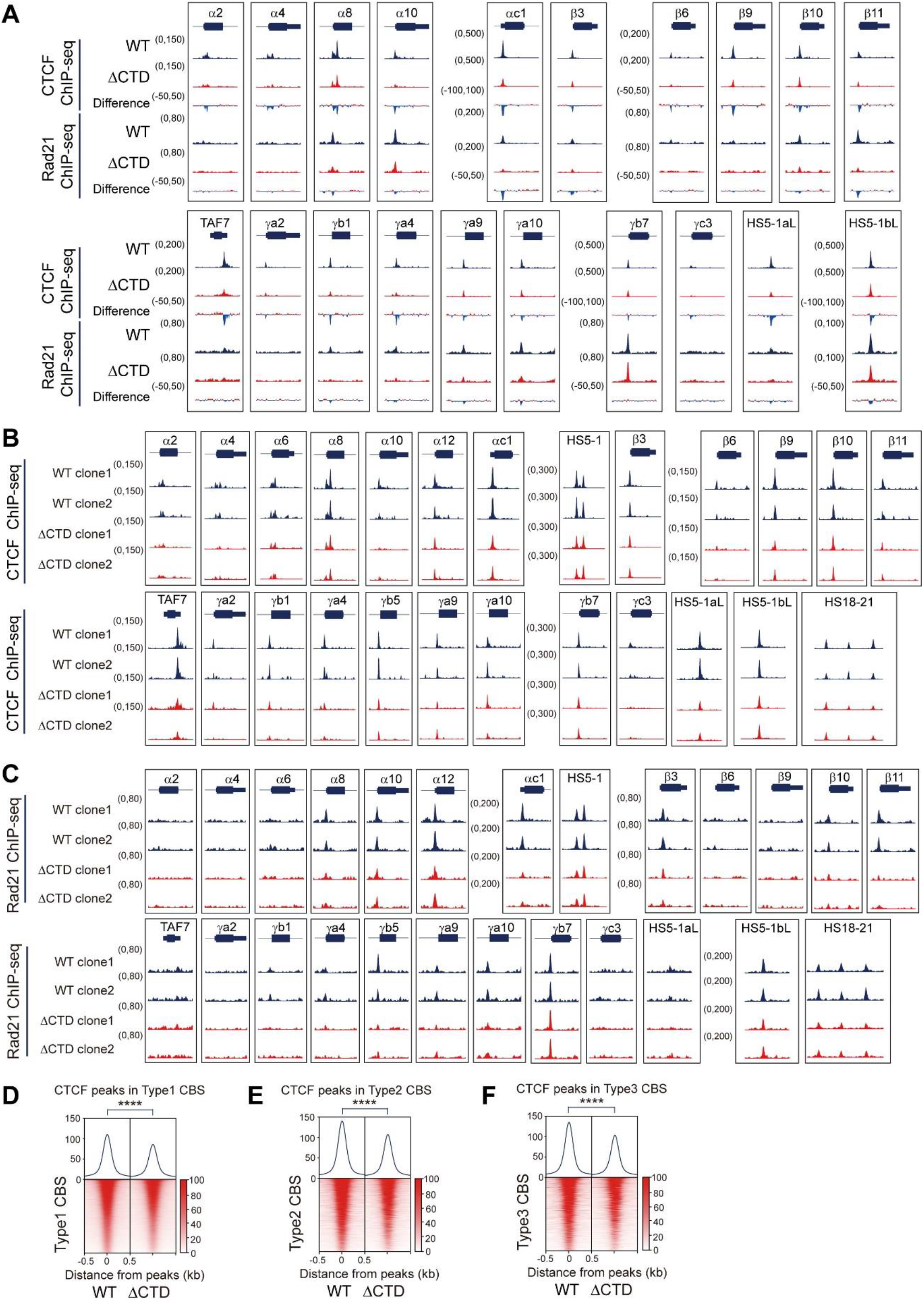
CTCF and Rad21 ChIP-seq profiles in ΔCTD and WT cells. Related to Figure 1. (A) CTCF and Rad21 ChIP-seq profiles for WT and ΔCTD cells at the *cPCDH* locus, indicating decreased CTCF binding upon CTD deletion. (B and C) CTCF (B) or Rad21 (C) ChIP-seq profiles for the two single-cell clones of WT or ΔCTD at the *cPCDH* locus. (D-F) Heatmaps of CTCF ChIP-seq signals at the three types of CTCF motifs upon CTD deletion. Student’s *t* test, \*\*\*\**p* < 0.0001.

**Figure S3.**
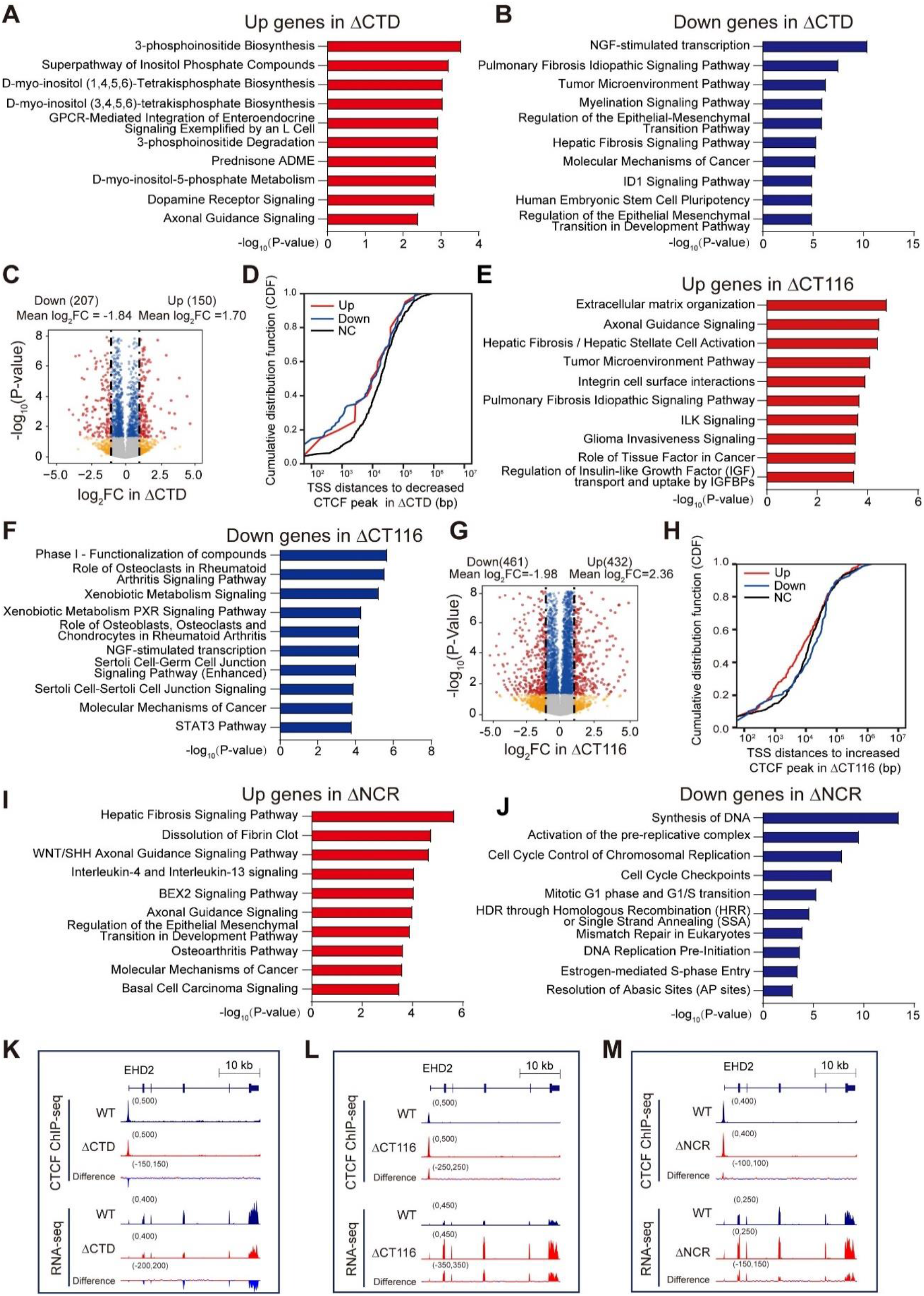
RNA-seq of ΔCTD, ΔCT116, ΔNCR cell clones. Related to Figures 1, 2, and 4. (A and B) Ingenuity pathway analysis (IPA) of up- (A) and down- (B) regulated genes upon CTD deletion. (C) Volcano plot of differential gene expression analyses upon ΔCTD deletion. Red dots, gene expression changed upon CTD deletion (log2FC > 1 and adjusted *p* value < 0.05). Blue dots, genes only passed adjusted *p* value < 0.05. Yellow dots, gene only passed log2FC >1. (D) TSS distances of up-, down-, or none-regulated (NC) genes to the closest decreased CTCF peaks in ΔCTD cells (data were shown as a cumulative distribution function, CDF). (E and F) IPA analyses of up- (E) and down- (F) regulated genes in ΔCT116 cells. (G) Volcano plot of differential gene expression analyses of WT and ΔCT116 cells. (H) TSS distances of up-, down-, or none-regulated (NC) genes to the closest increased CTCF peaks in ΔCT116 cells (data were shown as a cumulative distribution function, CDF). (I and J) IPA analyses of up- (I) and down- (J) regulated genes upon NCR deletion. (K-M) Altered CTCF binding and gene expression levels at the *EHD2* locus in ΔCTD (K), ΔCT116 (L), and ΔNCR (M) cell lines.

**Figure S4.**
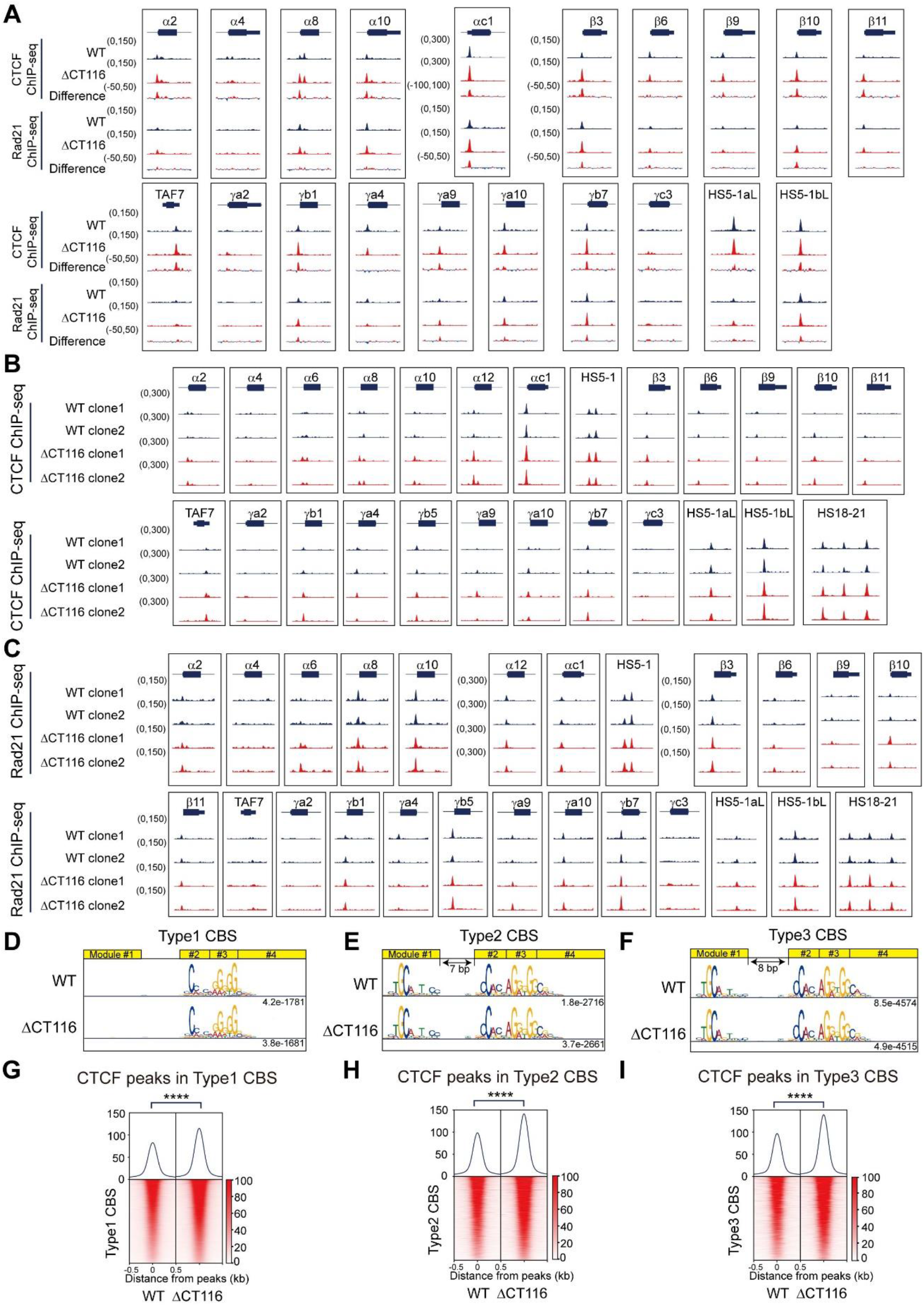
CTCF and Rad21 ChIP-seq profiles in ΔCT116 and WT cells. Related to Figure 2. (A) CTCF and Rad21 ChIP-seq profiles of WT and ΔCT116 cells at the *cPCDH* locus, showing increased CTCF binding in ΔCT116 cells. (B and C) CTCF (B) or Rad21 (C) ChIP-seq profiles of the two single-cell clones of WT and ΔCT116 at the *cPCDH* locus. (D-F) Three types of CTCF motifs of WT and ΔCT116 cells. (G-I) Heatmaps of CTCF ChIP-seq signals at the three types of CTCF motifs of WT and ΔCT116 cells. Student’s *t* test, \*\*\*\**p* < 0.0001.

**Figure S5.**
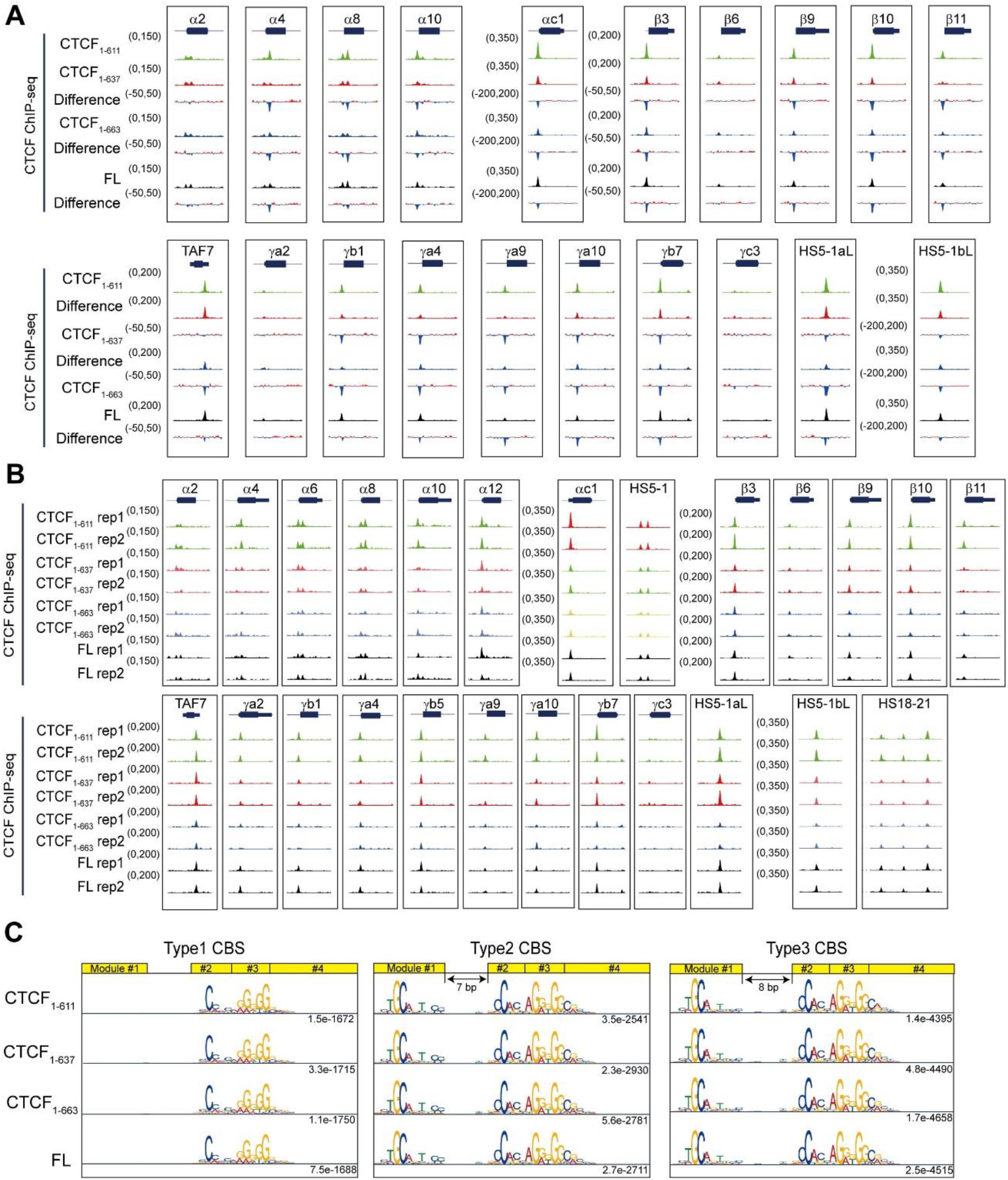
ChIP-seq profiles of a series of truncated CTCF proteins. Related to Figure 3. (A) ChIP-seq of different truncated CTCF proteins at the *cPCDH* locus, indicating that CTCF1-611 has the highest binding strength. (B) Biological replicates of ChIP-seq experiments with truncated CTCF proteins. (C) Motif analyses of truncated CTCF proteins, showing no alteration of the three types of CTCF motifs.

**Figure S6.**
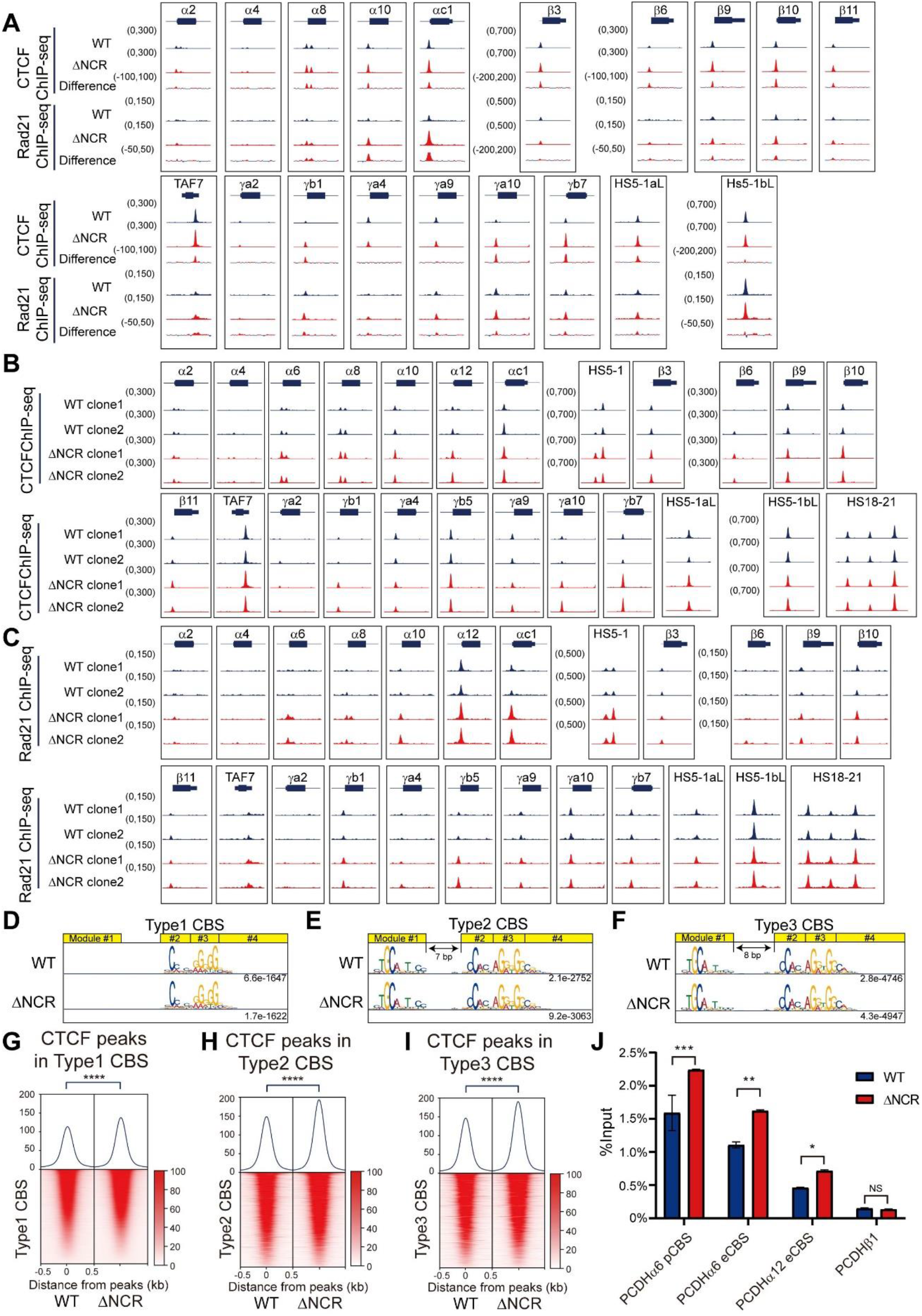
CTCF and Rad21 ChIP-seq profiles upon NCR deletion. Related to Figure 4. (A) CTCF and Rad21 ChIP-seq profiles at the *cPCDH* gene complex upon NCR deletion. (B and C) CTCF (B) or Rad21 (C) ChIP-seq profiles of the two single-cell clones of WT and ΔNCR at the *cPCDH* locus. (D-F) Three types of CTCF motifs upon NCR deletion. (G-I) Heatmaps of CTCF ChIP-seq signals at the three types of CTCF motifs upon NCR deletion. Student’s *t* test, \*\*\*\**p* < 0.0001. (J) CTCF ChIP-qPCR at three different CBS elements (*PCDHα6* pCBS, *PCDHα6* eCBS and *PCDHα12* eCBS) and a negative control (*PCDHβ1*) using WT and ΔNCR cell lines. Data are presented as mean ± SD. Student’s *t* test, \*\*\**p* < 0.001, \*\**p* < 0.01, NS, *p* > 0.05

**Figure S7.**
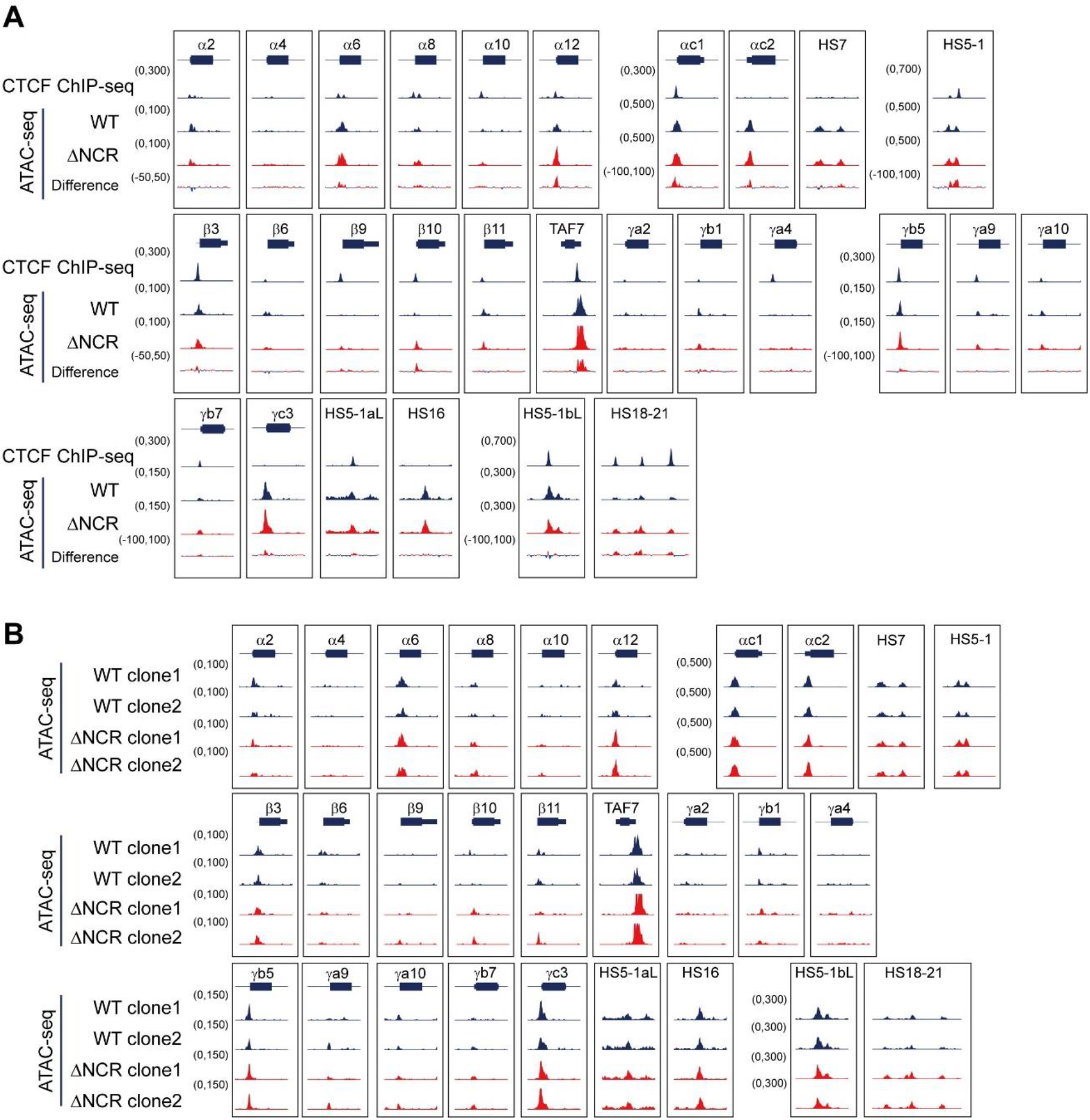
ATAC-seq profiles upon NCR deletion. Related to Figure 4. (A) ATAC-seq indicates increased chromatin accessibility at the *cPCDH* locus upon NCR deletion. (B) Close-up of ATAC-seq profiles at the *cPCDH* gene complex. (C) ATAC-seq profiles of the two single-cell clones of WT and ΔNCR at the *cPCDH* gene complex.

**Figure S8.**
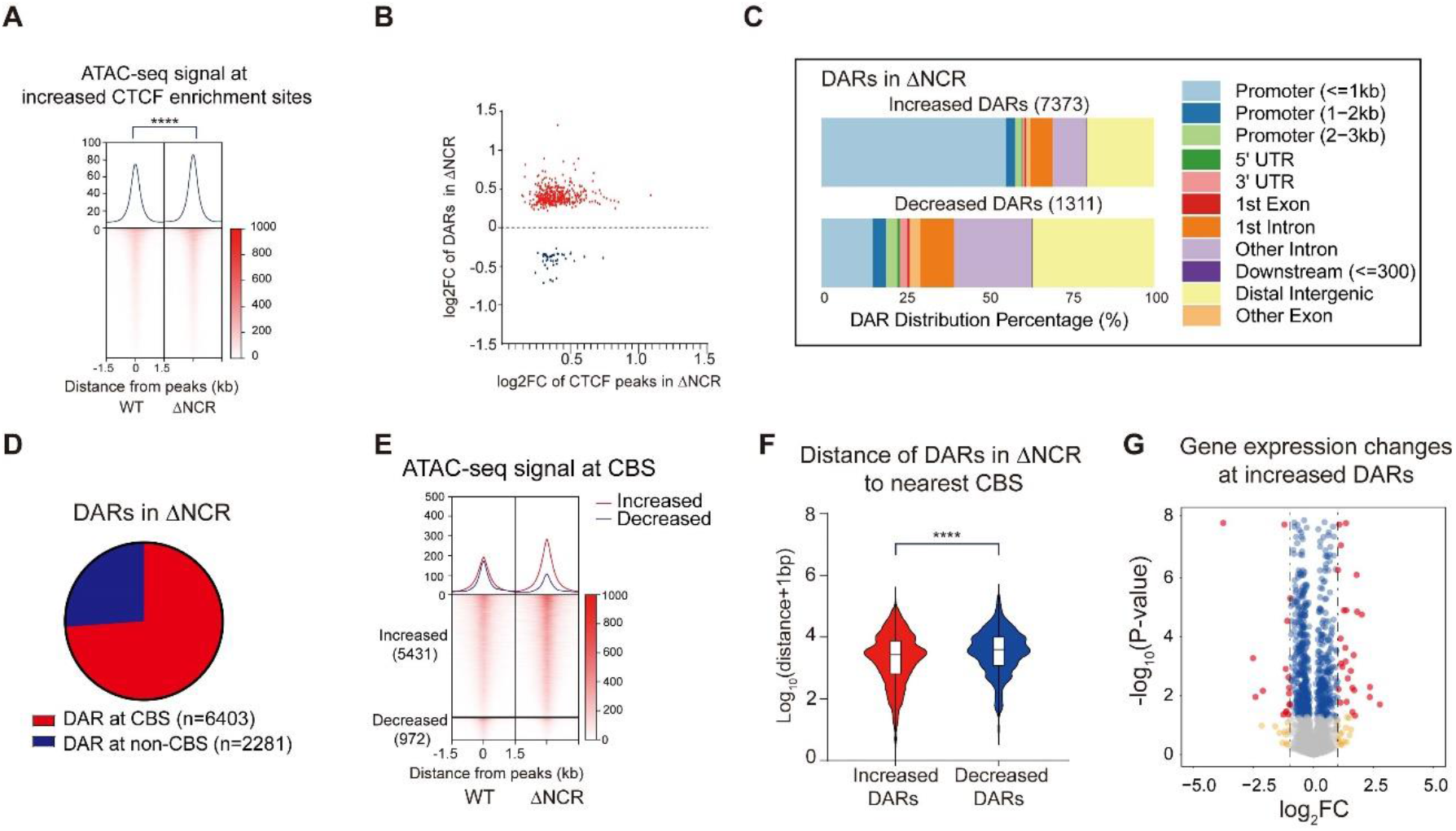
Deletion of NCR rewires chromatin accessibility. Related to Figure 4. (A) Heatmaps of ATAC-seq signals at sites with increased CTCF enrichments, indicating increased chromatin accessibility upon NCR deletion. Student’s *t* test, \*\*\*\**p* < 0.0001. (B) Scatter plot shows the log2FC of increased CTCF enrichments (x axis) and the corresponding log2FC of DARs (y axis) upon NCR deletion. (C) Distributions of increased and decreased DARs with gene elements upon NCR deletion. (D) Distribution of DARs in term of whether overlapping with CBS elements. (E) Heatmaps of DARs overlapped with CBS elements. (F) Violin plot of log10(distance+1bp) distribution of increased and decreased DARs to the nearest CBS element upon NCR deletion, showing that increased chromatin accessible regions are closer to CBS elements. Student’s *t* test, \*\*\*\**p* < 0.0001 (G) Volcano plot showing that increased DARs are associated with more up- regulated genes upon NCR deletion.

## Notes

### Competing Interest Statement

The authors have declared no competing interest.

### Summary of Updates

New Figure5 updated, Method updated. Supplementment Figure S1,S3,and S6 updated.

